# Sexually distinct song cultures in a songbird metapopulation

**DOI:** 10.1101/2021.07.05.451205

**Authors:** Wesley H. Webb, Michelle M. Roper, Matthew D. Pawley, Yukio Fukuzawa, Aaron M. Harmer, Dianne H. Brunton

## Abstract

Songbirds learn their songs culturally, through imitating tutors. The vocal culture of a songbird population changes as new song units (syllables) are introduced through immigration, copying errors, and innovation, while other syllables fall out of use. This leads to a diversification of the syllable pool across the species, much like the diversification and spatial patterns of human language. Vocal cultures have been well studied in male songbirds but have been largely overlooked in females. In particular, few studies compare spatial variation of male and female song cultures. Here we undertake one of the first comparisons of male and female song culture in birds, analysing song data from a metapopulation of New Zealand bellbirds *Anthornis melanura*, spanning an archipelago of six islands. Having classified 20,700 syllables, we compare population syllable repertoire sizes and overlap between sites and sexes. We show that males and females—both with complex songs—have distinct song cultures, sharing only 6–26% of syllable types within each site. Furthermore, male and female syllable types can be statistically discriminated based on acoustic properties. Despite diverse syllable repertoires within sites, few syllable types were shared between sites (both sexes had highly distinct site-specific dialects). For the few types shared between sites, sharing decreased with distance only for males. Overall, there was no significant difference between sexes in degree of site–site repertoire overlap. These results suggest different cultural processes at play for the two sexes, underlining the inadequacy of male-centric song research and calling for comparisons of male and female song cultures in many more species.

## Introduction

Culture is shared information or behaviour acquired through social learning from conspecifics (Dawkins, 1976), involving the transmission of *memes* (units of culture) by behavioural imitation. Vocal culture—the social learning of acoustic memes—has so far been observed in songbirds (oscines; Passeri), some suboscines (*Procnias* spp., Cotingidae), parrots, hummingbirds, cetaceans, elephants, seals, bats, and humans (Paton et al., 1981; Baptista and Schuchmann, 1990; Janik and Slater, 1997; Poole et al., 2005; Sanvito et al., 2007; Catchpole and Slater, 2008; Kroodsma et al., 2013). In these taxa, the vocal repertoire of a population changes as new memes are introduced through immigration, copying errors, and innovation, while other memes fall out of use (Catchpole and Slater, 2008). This leads to a diversification of the meme pool across the species, much like the diversification of human language—resulting in dialects (Podos and Warren, 2007).

Despite the high volume of studies on male birdsong culture and dialects (Jenkins, 1978; Whitehead and Rendell, 2014; Aplin, 2019), little is known about female song culture. This is partly due to a northern-hemisphere-biased view of sexual selection that emphasises male-male competition and female choice in driving elaborate traits in males (Darwin, 1871), but which does not provide a framework for understanding elaborate female traits (Riebel et al., 2019). Female song (and other elaborate female traits) have been overlooked as non-functional aberrations, resulting ‘accidentally’ from shared genetic architecture with males (Darwin, 1871; Lande, 1980; see Tobias et al., 2012 for review). This view has now been roundly discounted. Female song is present in 64% of surveyed songbird species (Webb et al., 2016), has been recovered as the ancestral state (Odom et al., 2014), can evolve independently of the male song phenotype (Price, 2015) and has female-specific functions in territory and resource defence, mate attraction, mate defence, and pair bonding (reviewed in Austin et al., 2021).

Overlooking females has impeded development of a more general theory that explains song culture in both sexes. For instance, the possibility that females have song cultures distinct from conspecific males, with different geographic patterns of song sharing, has hardly been investigated at a population level (Graham et al., 2017a).

Studies have reported a wide degree of sexual song dimorphism in songbirds, from identical repertoires for males and females at one extreme [e.g., forest weaver *Symplectes bicolor* (Wickler and Seibt, 1980), magpie lark *Grallina cyanoleuca* (Hall, 2000)], to completely non-overlapping repertoires at the other [e.g., many duetting wrens; (Brown and Lemon, 1979; Levin, 1996; Mann et al., 2009)]. However, studies comparing the sexes have mostly focused on within-pair repertoires in duetting species. To our knowledge, only four studies (of just two species) have compared spatial variation of male and female songs at a population level.

First, Mennill and Rogers (2006) examined the songs of duetting male and female eastern whipbird *Psophodes olivaceus* across their geographic range. While male whipbird song was highly consistent over space, female song showed pronounced variation, with multiple distinct song types. The authors suggest eastern whipbirds have undergone a decoupling of male and female song learning strategies in response to different sexspecific selection pressures.

Three later studies on rufous-and-white wrens (Graham et al., 2017a, 2018a, 2021) showed that song cultures of males and females can be similar in complexity, and appear to evolve in similar directions via acoustic adaptation and cultural drift. However, males and females differ in the relationship between dispersal distance and song-sharing with parents (Graham et al., 2017a), the speed of cultural change, and the relationship between immigration rate and cultural diversity (Graham et al., 2021).

We know of no other studies to examine spatial variation in song culture at a population level, comparing both sexes.

The New Zealand bellbird *Anthornis melanura*—hereafter ‘bellbird’—provides an ideal system for comparing male and female song cultures over space. The bellbird is a non-duetting endemic honeyeater (Family Meliphagidae) with complex, geographically diverse song in both sexes, and probable open-ended learning (Roper, 2018). Bellbird populations occur across a network of islands and peninsulas in the Hauraki Gulf, northeastern New Zealand. Population connectivity is substantial, and regulated by geographic isolation (Baillie, 2011; Baillie et al., 2014). Females disperse more frequently than males, resulting in higher female connectivity between sites (as typical for songbirds: Greenwood, 1980; Clarke et al., 1997; Paris et al., 2016).

In this paper, we seek to advance understanding of female birdsong culture by comparing the patterns of male and female meme sharing across a network of six island and peninsula populations (Figure 1). Our questions are:

1. *Are there sexual differences in population syllable repertoire size?*
2. *How much do male and female population repertoires overlap?*
3. *What is the pattern of syllable type sharing between sites?*

**Figure 1.**
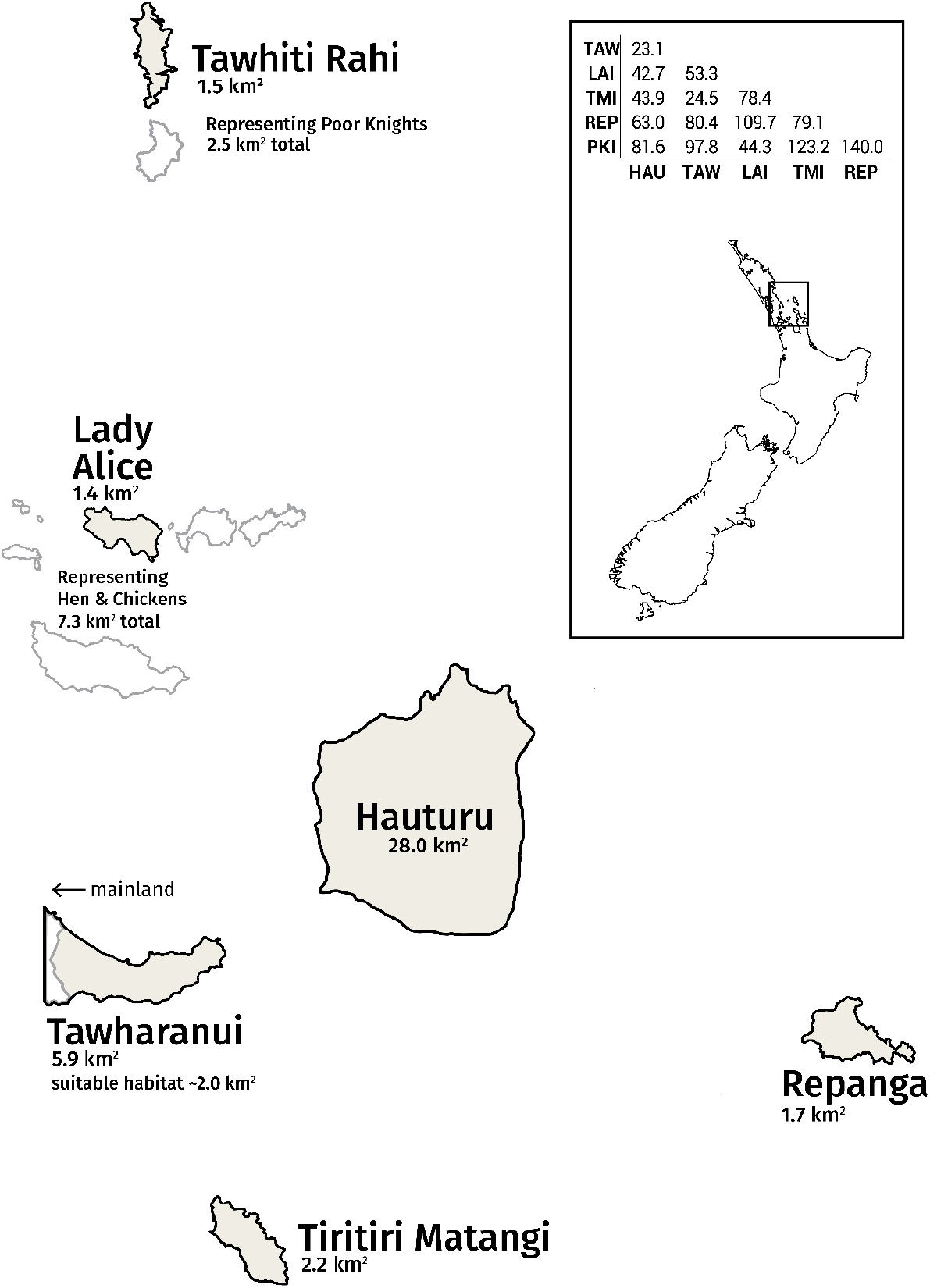
Simplified map of the Hauraki Gulf archipelago, home to a metapopulation of bellbirds. Tawhiti Rahi represents the Poor Knights Islands group; likewise, Lady Alice Island represents the Hen and Chickens Islands group. The other islands of these wider groups are shown in white. The distance matrix between sites (measured in km) is provided (HAU, Hauturu; TAW, Tawharanui; LAI, Lady Alice; TMI, Tiritiri Matangi; REP, Repanga, PKI, Tawhiti Rahi). Distances are not to scale on simplified map.

If geographic proximity drives cultural connectivity between islands, we expect sites that are closer together will share more memes (i.e., an isolation-by-distance pattern in vocal culture). We also expect higher female dispersal frequency will result in higher meme flow between sites, and thus higher sharing of syllable types between sites than for males.

## Methods

### Creating the song database

#### Recording bellbird song

Recording trips were conducted to six island and mainland peninsula sites in the Hauraki Gulf 2013-2017 (Table 1), with iwi consent (Ngatiwai, Ngati Manuhiri) and permits from the New Zealand Department of Conservation (47948–FAU, 34833–FAU, 41756–FAU, 48000–FAU) and Massey University Animal Ethics Committee (permit number 15/21). The sites were chosen based on their large bellbird populations (good return for sampling effort) and their variety in connectivity. A team of 1–5 recordists per site collectively spent a total of over 1,000 recordist-hours actively tracking and recording wild bellbirds (per site: Tawhiti Rahi, 108; Lady Alice, 110; Hauturu, 140; Tawharanui, 132; Repanga, 260; Tiri, many hundreds of hours), with Marantz PMD661 portable solid-state recorders paired with handheld Sennheiser ME-66 shotgun microphones. Recordists coordinated movement to sample sites systematically and with maximal coverage, gathering 2,137 high-quality recorded songs (discrete vocalisation bouts) by adult individuals of known sex, during daylight hours. During recording, metadata including identity, age and sex of the focal bird was spoken into the microphone for later transcription.

**Table 1—.**
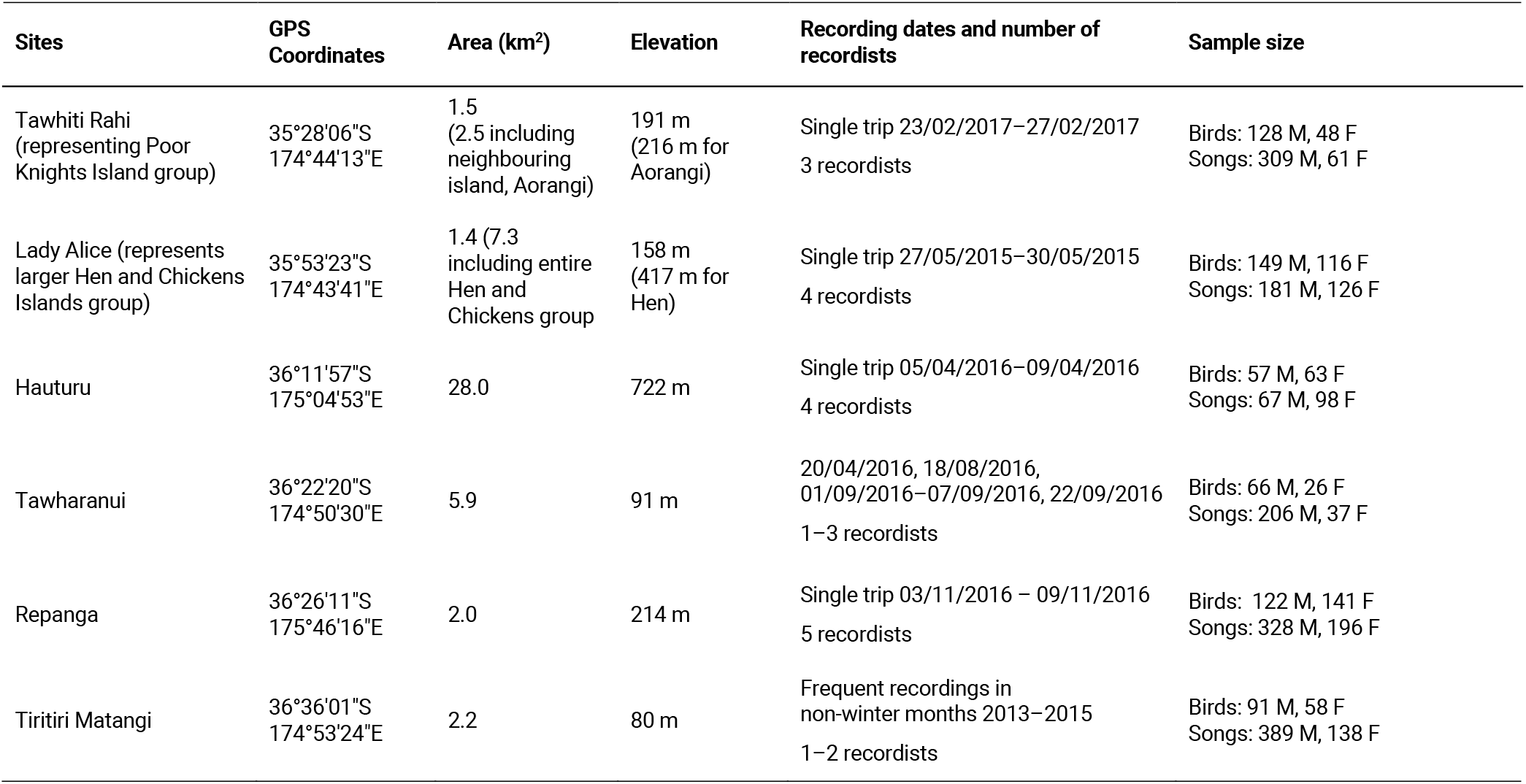
Geographic and sampling information for the six study sites. Tawhiti Rahi represents the larger Poor Knights Islands group; likewise, Lady Alice Island represents the larger Hen & Chickens Islands group. Elevation data is from www.topomap.co.nz. In the *Sample size* column, M=male and F=female. Recording on Hauturu was restricted to an area ~3 km^2^.

#### Segmenting songs into syllables

Songs were imported into a database in *Koe* bioacoustics software (Fukuzawa et al., 2020) and manually segmented into syllables by visually inspecting the spectrogram and setting syllable start/end points. Bellbirds mostly sing in discrete units that are easy to demarcate as syllables based on a gap either side. Occasionally there are fast-paced bursts where it is not clear where one unit ends and another begins; in these cases sounds were grouped together as a syllable if they were separated by less than 15 ms of silence. This value was chosen through trial and error for the best consistency of syllable groupings.

#### Classifying syllables

To prepare for classification, the 20,700 segmented syllables were ranked by acoustic similarity in *Koe*. Similarity was calculated by extracting all available acoustic features for each unit and applying the UPGMA method (Sokal, 1958); see https://github.com/fzyukio/koe/wiki for details of extracted features and similarity index calculation.

WHW then compared syllables visually and aurally to classify them into types based on just-noticeable differences in pitch, timbre, and duration of the playback and visual appearance of the spectrogram. Manual perception-based classification is considered excellent for acoustic classification (Sayigh et al., 2007; Duda et al., 2012) and allowed finer scale than automated classification methods, such as multi-dimensional scaling (MDS) or principal component analysis (PCA). The high acuity of bird hearing relative to humans (Dooling, 2004) justifies a fine-scale classification approach. We validated our classification by having 74 inexperienced judges independently label a subset of syllables. Average match with our own labels was 89.6% (median 95.6%); see Fukuzawa et al. (2020) and Article S1A for more details.

### Sex differences in population-level repertoires

#### Male and female syllable diversity

The raw number of syllable types recorded at each site are not directly comparable, due to the inevitable confounding of different sampling effort. To account for this, we used the statistical software *EstimateS* (Colwell, 2013) to produce syllable type accumulation curves and extrapolate the true number of syllable types at each site (Article S1B).

#### Repertoire overlap of male and female population sectors

For each site we calculated the percentage overlap between male and female population repertoires, using the Jaccard similarity index (Hamers et al., 1989): the number of shared types divided by the total number of types.

Next, the acoustic feature measurements of all syllables (previously extracted to aid classification in *Koe*) were normalised by mean-centering and dividing by standard deviation. To test whether male-only, female-only, and shared syllable types form three distinguishable clusters in acoustic space, we classified syllables using a linear discriminant analysis (LDA) on the data, with leave-one-out cross-validation. If male-only, female-only, and shared syllables are randomly interspersed in acoustic space, then the LDA classification results will be similar to random assignment of groups. If the three groups occupy more distinct ranges, then the LDA classification will perform much better than random assignment.

### The pattern of syllable type sharing between sites

We calculated repertoire overlaps between sites, for all pairwise combinations of site lists—comparing males against males, females against females. For each sex we tested for an isolation-by-distance pattern in repertoire overlaps between sites, in two ways.

First, we conducted a Hauturu-centric analysis. The large size, geographic centrality, and the long-term persistence of the Hauturu bellbird population make it likely to be a substantial source of dispersing bellbirds to other sites in the Hauraki Gulf archipelago. We calculated Spearman’s rank correlation coefficients between *population repertoire overlap with Hauturu* against *geographic distance from Hauturu*.

Next, to test for an overall isolation-by-distance pattern, we calculated Spearman’s rank correlation coefficients between *percentage of site–site repertoire overlap* against *site–site geographic distance*, for all pairwise combinations of sites in the archipelago. In this case, to account for non-independence of points (being pairwise combinations), we used the RELATE routine in *PRIMER* (version 7, Clarke and Gorley, 2015) with 9999 permutations. To test whether males or females share more syllables between sites overall, we used a two-tailed Sign Test (Dixon and Mood, 1946), with null hypothesis: the degree of site–site sharing is equal between sexes (Article S1C).

## Results

### Sex differences in population repertoires

#### Male and female syllable diversity

Across the metapopulation we recorded a total of 702 syllable types: 337 male-only, 203 female-only, and 162 shared. See Data S1 and Table S1 for a complete catalogue.

For each site and sex, the syllable type accumulation curve levelled off nearly or completely by 500 songs (i.e., approximating the true number of syllable types present), allowing meaningful comparison of syllable diversity (Figure S1).

We found no evidence of a difference in repertoire size for males versus females within sites; 95% confidence intervals overlap for males and females in all cases (Table 2). There was high variability between sites for both sexes, with male point estimates ranging from 80 (Repanga) to 186 (Hauturu) and females from 81 (Repanga) to 186 (Hauturu).

**Table 2—.** Extrapolated estimates of male and female syllable diversity within sites. Estimated number of syllable types at 500 songs, for male and female population sectors at each site. Ninety-five-percent confidence intervals are in brackets. Calculated with the statistical software *Estimate S* (Colwell, 2013).

| Site | Male | Female |
| --- | --- | --- |
| Tawhiti Rahi | **135** (121–149)<br>*n*=306 songs | **119** (68–169)<br>*n*=58 songs |
| Lady Alice | **143** (118–169)<br>*n*=181 | **128** (93–163)<br>*n*=125 |
| Hauturu | **186** (139–234)<br>*n*=67 | **186** (151–220)<br>*n*=98 |
| Tawharanui | **116** (103–129)<br>*n*=204 | **89** (48–130)<br>*n*=37 |
| Repanga | **80** (72–89)<br>*n*=328 | **81** (65–97)<br>*n*=196 |
| Tiri 2013, 2014, 2015 | **165** (152–178)<br>*n*=389 | **145** (115–175)<br>*n*=138 |

#### Repertoire overlap of male and female population sectors

Male and female population sectors had largely separate repertoires, ranging from 6% total units shared at Tawhiti Rahi to 26% shared at Hauturu (Figure 2, Table S2). In other words, 74-94% of syllable types at each site were sung by one sex only.

**Figure 2.**
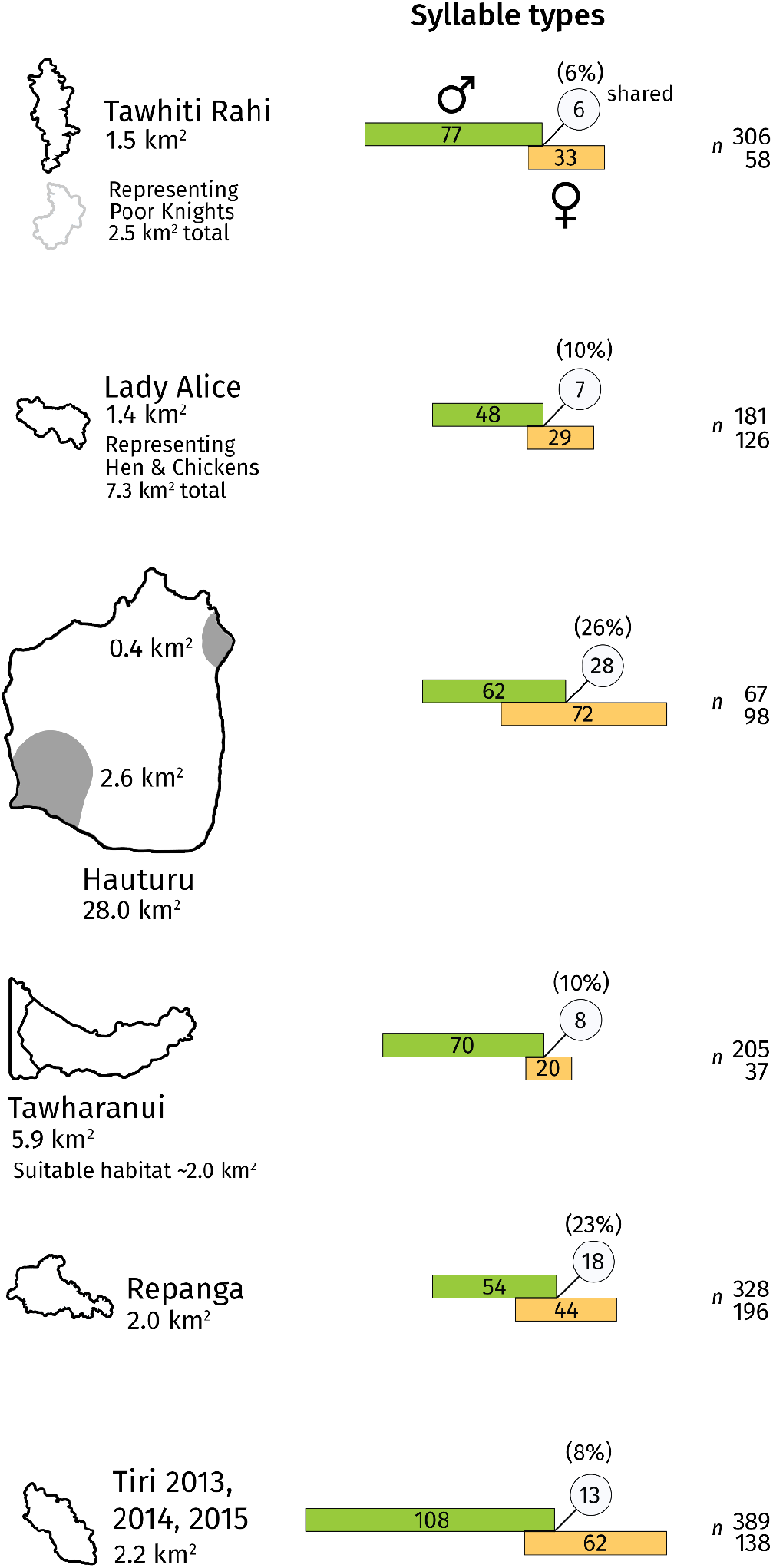
Overlap of male and female population syllable repertoires. Each diagram indicates the recorded number of male (green bar) and female (orange bar) syllable types, and the number of types common to both sexes (circle). The percentage overlap is given above the circle, calculated as the Jaccard similarity index: number of shared types divided by total number of types. For robustness, repertoires exclude types with fewer than three occurrences within that site-and-sex group. Bar length is proportional to the number of syllable types, and overlap length is proportional to the number of shared syllable types. Sample sizes (number of songs) for the male and female bars are given to the right of the bars. Note that repertoire sizes are recorded values, not extrapolated, and therefore are not directly comparable due to differing sample sizes. However, relative overlap percentage is more robust to differing sample sizes and thus meaningful to compare. Site outlines on the left of the figure are to scale. The shaded grey area in the Hauturu outline indicates the sampled region of the island.

**Figure 3.**
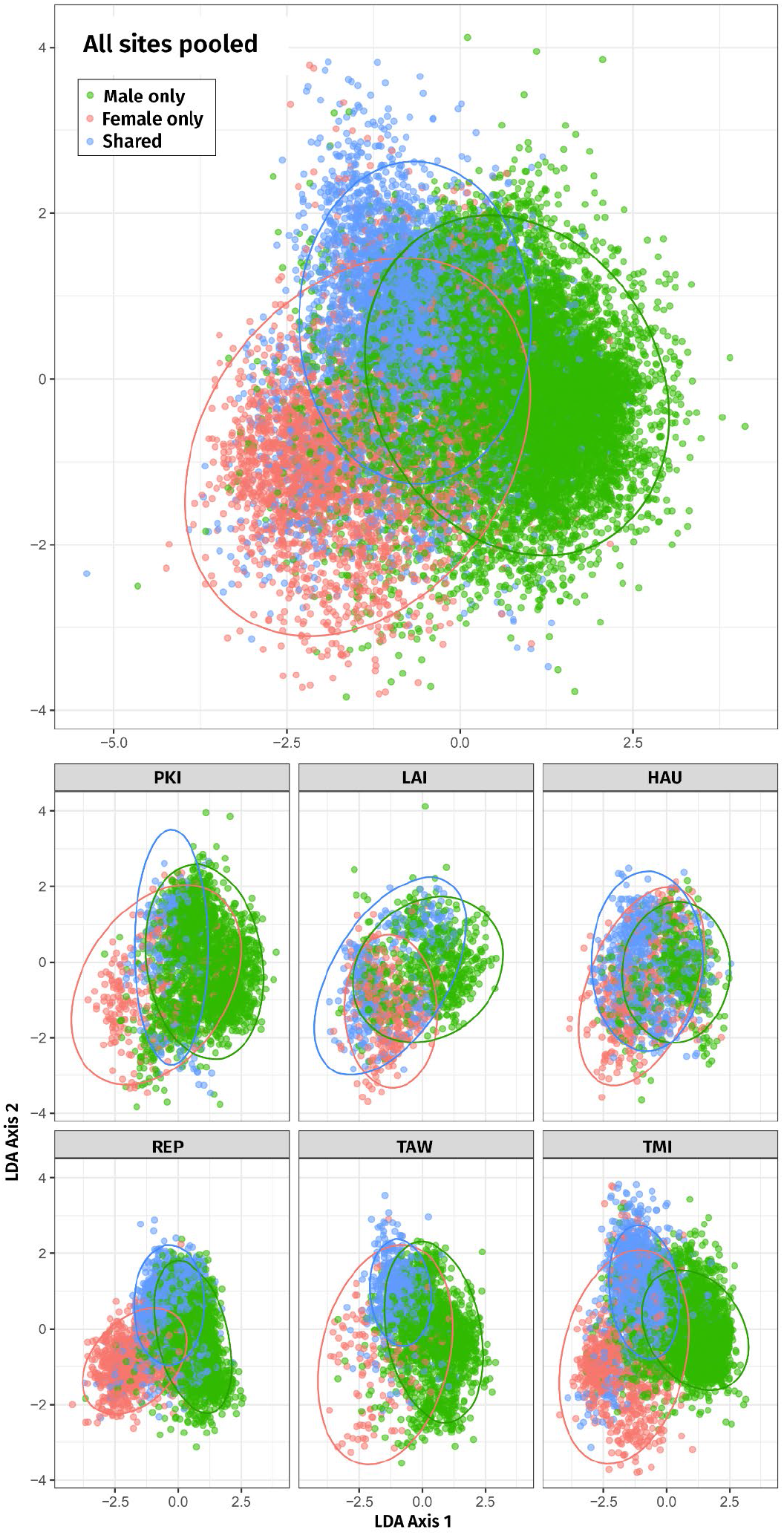
Two-dimensional linear discriminant analysis (LDA) of male-only, female-only, and shared syllables for all sites combined (top) and each site separately (bottom). Ellipses represent the regions covering 95% of the data (according to the fitted normal). Each site is plotted using the same axes to be comparable. Site abbreviations are as follows: PKI, Tawhiti Rahi, representing Poor Knights Islands; LAI, Lady Alice Island, representing Hen & Chickens; HAU, Hauturu; REP, Repanga; TAW, Tawharanui; TMI, Tiri.

In the Linear Discriminant Analysis (LDA) ordination, male-only and female-only syllables form two clear, partially-overlapping clusters, with shared syllables in between, when data from all sites are pooled. For individual sites the configuration of the three clusters varies but shows a tendency for female-only syllables to occur at lower LD1 values, male-only to occur at higher LD1 values, with shared syllables in between.

Success of the LDA leave-one-out classifier had an overall *lift* of 1.78; that is, the LDA assigned labels (male-only, female-only and shared) to syllables overall 1.78 times better than random allocation of labels. Lift was 1.51 for male-only syllables, 3.99 for female-only syllables, and 2.15 for shared syllables. P values for all lift tests were <0.001; i.e., in all cases none of the 4999 null distributions had a lift as large as the observed lift. Therefore, male-only, female-only, and shared types can be separated on the basis of acoustic features.

### The pattern of meme sharing between sites

#### Site–site repertoire overlap

Matrices of site–site repertoire overlap are presented in Figure 4. The degree of site-site repertoire overlap between male populations was low, ranging 3-18% (median=6%); for females: 1-12% (median=5%). Remarkably, for both sexes, the syllable types shared between sites were almost exclusively pure-tone whistle syllables or simple stutter-like syllables. The one exception was Hauturu–Tawharanui, which additionally shared some more complex types following the recent Hauturu→Tawharanui founder event (Brunton et al., 2008). See Data S1 and Data S2.

**Figure 4.**
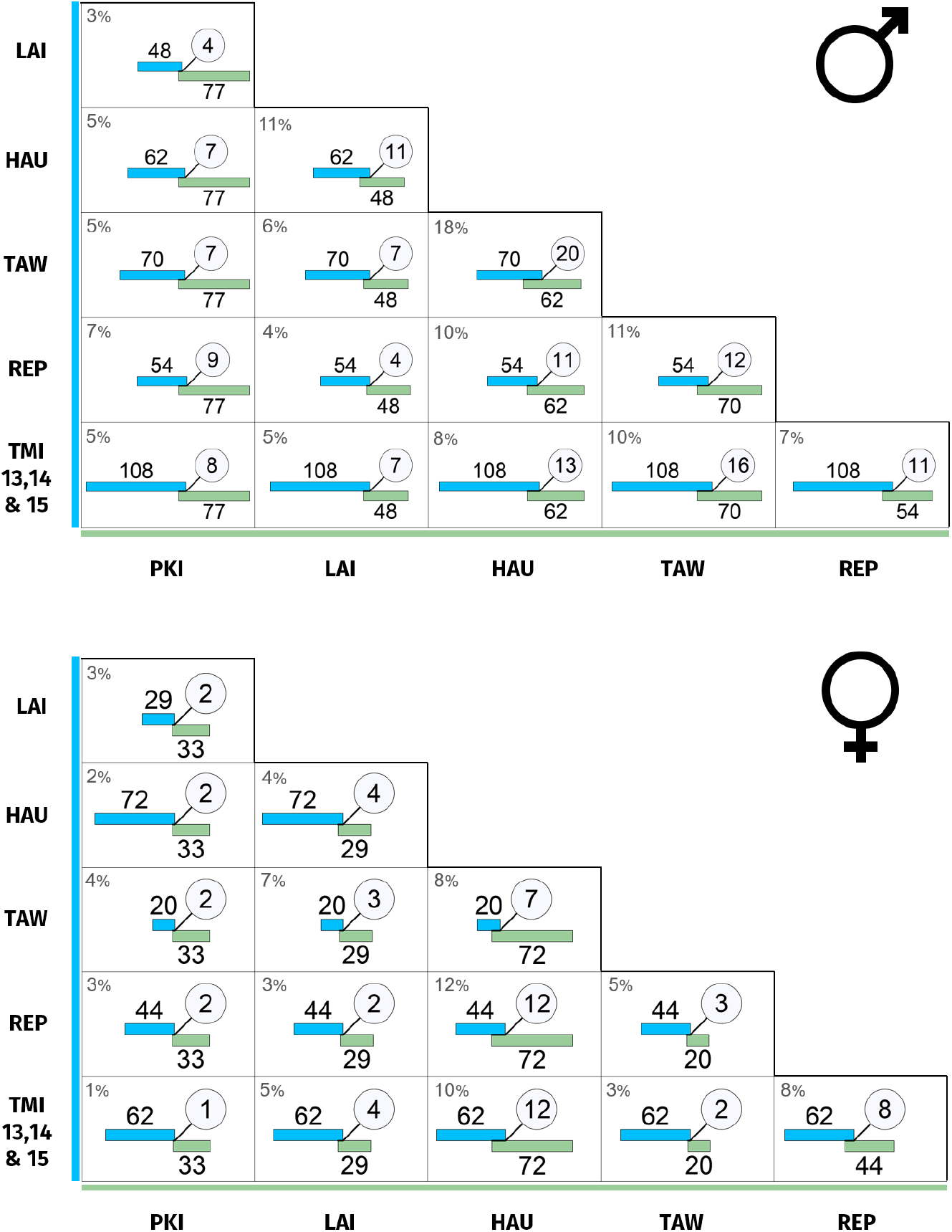
(*Top*) Male population repertoire overlap between sites. (*Bottom*) Female population repertoire overlap between sites. Each cell indicates the recorded number of syllable types for the two sites (blue and green bars), and the number of types common to both (circle). The percentage overlap is given at the top left of each cell and is calculated as Jaccard similarity index: number of shared types divided by total number of types. For robustness, repertoires exclude types with fewer than three occurrences within that site and sex. Bar length is proportional to the number of syllable types, and overlap length is proportional to the shared number of syllable types. Site abbreviations are as follows: PKI, Tawhiti Rahi (representing Poor Knights Islands); LAI, Lady Alice Island (representing Hen and Chickens); HAU, Hauturu; TAW, Tawharanui; REP, Repanga; TMI, Tiri.

#### Repertoire overlap versus distance

The relationship between repertoire overlap with Hauturu versus geographic distance from Hauturu is shown in Figure 5A. There was a negative correlation between sharing and distance for males (rs = −0.90, *df*=3, 0.03<P<0.05) but no relationship for females (rs =-0.10, *df*=3, P>0.10). Contrary to predictions, sites at increasing distances from Hauturu did not share a progressively diminishing subset of syllable types with Hauturu, but different (apparently unrelated) subsets; this was true for both males and females (Data S2).

**Figure 5.**
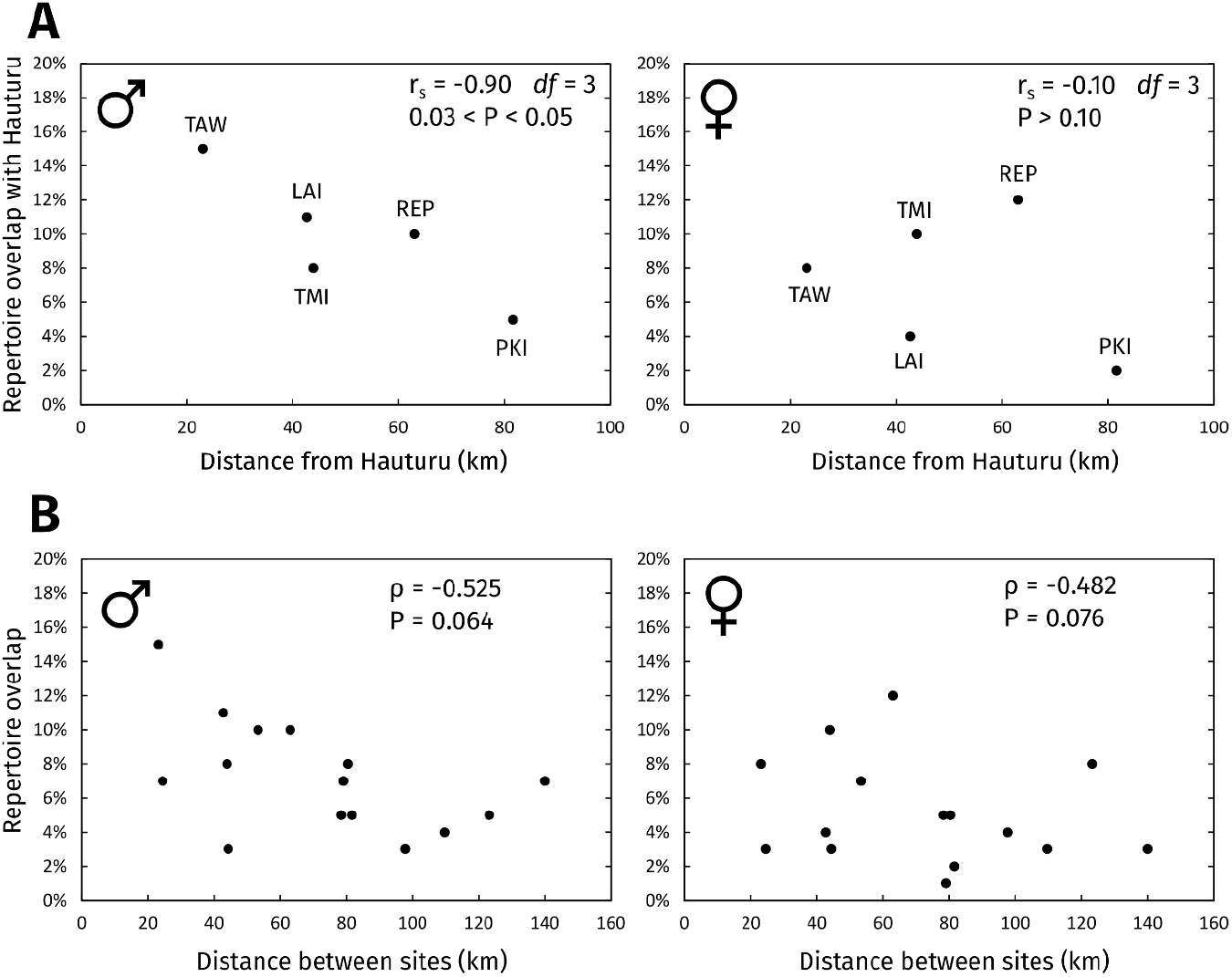
Repertoire overlap versus distance between sites. **(A)** Repertoire overlap of all subpopulations with Hauturu versus their distance from Hauturu. Overlaps were calculated on repertoire lists of types with 3+ occurrences within each site-and-sex group. Site abbreviations: TAW, Tawharanui; LAI, Lady Alice Island (representing Hen and Chickens); TMI, Tiri; REP, Repanga; PKI, Tawhiti Rahi (representing Poor Knights Islands). **(B)** Site-site repertoire overlap versus site-site geographic distance, for all pairs of sites. Spearman’s rank correlation coefficients (rs) were calculated using the RELATE routine in *PRIMER* (version 7) with 9999 permutations, accounting for the non-independence of points.

When considering sharing between *all* pairwise combinations of sites in the archipelago (Figure 5B), we did not find evidence of an isolation-by-distance pattern, for males (ρ= −0.525, P=0.064) or females (ρ=-0.525, P=0.076).

The recent founder event, where bellbirds from Hauturu colonised Tawharanui (Brunton et al., 2008) could possibly inflate the degree of sharing between the two sites, compared to other sites where both populations are well established. Therefore we tested robustness of the correlations to removal of the Hauturu–Tawharanui datapoint, but found little change in the slope or strength of the correlations.

There was no evidence of an overall difference between males and females in the degree of site-site repertoire overlap; out of 15 site-site comparisons, males shared a higher percentage of syllables than females in 9 cases, lower in 4 cases, and equal in two cases (Sign Test P=0.17).

## Discussion

This is one of the first comparisons of male and female song cultures across a metapopulation (Mennill and Rogers, 2006; Graham et al., 2018a, 2018b, 2021). We found that male and female bellbirds have comparable repertoire sizes (at a population level; Table 2), and sexually distinct vocal cultures, sharing only a small percentage of syllable types between sexes (6-26% within each site; Figure 2). Whether a type is male-specific, female-specific, or shared between sexes can be predicted based on its acoustic properties (Figure 3). Furthermore, song cultures of both sexes differ dramatically between sites— demonstrating male and female song dialects (Figure 4, Data S1). Despite a large and varied repertoire *within* sites, generally the only types shared *between* sites were flat-contour, pure-tone whistle syllables (Article S1A) or simple stutter-like syllables. Between-site sharing of these syllable types decreased with distance for males but not females. These contrasting patterns of sharing across the archipelago may result from sex differences in dispersal, meme mutation rates and song-learning modes.

Comparisons of male and female repertoire sizes typically focus on individual-level repertoires. Such studies have found smaller female repertoires in some cases (e.g., rufous-and-white wrens *Thryophilus rufalbus*, Mennill and Vehrencamp, 2005; banded wrens *Thryophilus pleurostictus*, Hall et al., 2015), equal-sized repertoires in other cases (e.g., bay wren *Thryothorus nigricapillus*, Levin, 1996), and at least one case of larger female repertoire size (stripe-headed sparrow *Peucaea r. ruficauda*, Illes, 2014). The drivers of individual repertoire size are thought to differ between sexes, with males under selection from male-male competition and female choice (Catchpole, 1987; Hill et al., 2018), and females perhaps primarily from female-female competition for non-sexual breeding resources (Tobias et al., 2012). In the present study, similar male and female repertoire sizes may be explained by bellbird social ecology. Bellbirds are socially monogamous, and both sexes are highly social and aggressive in singing interactions (Roper, 2018). It is plausible, therefore, that similar intensity of competition and social interaction drive the evolution of similar syllable diversity in the two sexes. However, our analysis is on population-level repertoires, which are a product of both individual repertoire sizes and variation between individuals. For example, equal male and female population repertoire sizes could also result if male individual repertoires are larger and more consistent between individuals (cultural conformity; Aplin et al., 2015), and females smaller and more variable (cultural non-conformity; Riebel et al., 2015). Additional focused recording of banded individuals is required to quantify these two sources of diversity.

It is interesting that population repertoire sizes varied greatly between sites. Adaptation to differing acoustic environments (Potvin and Clegg, 2015; Graham et al., 2017b) seems an unlikely explanation, as all sites were coastal, with similar vegetation (though soundscape was not measured). Another potential explanation is that genetic diversity has driven population repertoire size. However, sites with low genetic diversity (Baillie, 2011) did not have correspondingly low syllable diversity, suggesting song culture is not tightly constrained by genetic diversity in bellbirds (see also Graham et al., 2018). We suspect that between-site differences in repertoire size are more likely driven by competition level. For example, sites with higher population density may have elevated competition for food, or sites with high population connectivity may experience increased competitive encounters with migrants—selecting for bigger individual repertoires.

The discovery of sexually distinct, yet partly overlapping syllable repertoires raises questions about transmission and function. Logically, all sex-specific syllable types must be learnt male-to-male or female-to-female (as in rufous-and-white wrens, for example; Mennill and Vehrencamp, 2005). But what of types common to both sexes? These must reflect inter-sexual learning in some form, whether accidental or intentional (Evans and Kleindorfer, 2016). Once learnt inter-sexually, these memes could be transmitted intra-sexually and become an established part of the repertoire for that sex. Roper et al. (2018) found that juvenile male and female bellbird song is spectrally similar, then diverges prior to crystallisation. Perhaps both sexes of bellbird are physiologically capable of overlapping in acoustic space, but other factors (e.g., learning strategies, sexual/social selection) prevent it in the wild. This appears true in slate-coloured boubous *Laniarius funebris*; male and female boubous share no syllable types in the wild, but birds hand-raised under experimental conditions develop syllables of both sexes (Wickler and Seibt, 1988). Wild birds may choose to express sex-specific memes to help avoid being mistaken for the other sex, which risks attracting same-sex rivals or repelling potential mates (Logue et al., 2007). However, the benefit of *shared* syllables is unclear. Might sex-specific syllables function in intra-sexual communication, and shared syllables in inter-sexual communication? Sophisticated field experiments with banded individuals are needed to resolve meme functions and transmission modes in bellbirds.

We found that male-specific and female-specific syllables do occupy two largely distinct regions of acoustic space, with shared syllables occupying a cluster between in a ‘sex-neutral’ range (Figure 3). In the ordination, the separation of male-specific and female-specific memes could be due to universal morphological constraints of body size and syrinx structure, as males are 20% larger than females (Heather and Robertson, 2000) and have different syrinx morphologies (Roper, 2018). At the same time, the spread and shape of the clusters vary widely between sites, which may reflect site-specific cultures.

The pattern of meme sharing between sites defied our expectations. Repertoire similarity was not strongly related to geographic proximity between islands; only males showed some evidence of isolation with distance from Hauturu (Figure 5). However, for both sexes the percentage of syllable types shared between sites was small, and almost totally limited to pure-tone whistle syllables and simple stutter-like syllables (except for Hauturu-Tawharanui, which shared more types, likely due to the recent founder event described in Brunton et al., 2008). The observed pattern suggests that immigrants abandon most source memes after arrival, retaining only pure-tone and stutter-like types. Perhaps all but these simple types incur high aggression at new sites (‘colony password’ hypothesis; Feekes, 1977) and so are dropped in favour of the local dialect.

The large size and geographic centrality of the Hauturu population make it a likely source of dispersing bellbirds to other sites in the archipelago. Thus, one might expect the islands around Hauturu to form a chain of ‘stepping stones’ for bellbird dispersal, leading to progressively more dissimilar repertoires away from the central source population. There are many examples of such chains (Irwin et al., 2005; Parker et al., 2012; Lachlan et al., 2013). In contrast, we found that for both male and female bellbirds, sites at increasing distances from Hauturu did not share a progressively diminishing subset of syllable types with Hauturu, but different (apparently unrelated) subsets (Data S2). Therefore, our results suggest direct dispersal to each site from Hauturu, rather than serial dispersal along the island chain.

We predicted that female-biased dispersal in bellbirds should result in higher female meme flow and thus higher inter-site sharing for females than for males. Instead, we found no evidence for a sexual difference in amount of sharing between sites. That females do not show higher inter-site sharing suggests that other processes—such as a higher turnover rate or weaker retention of source memes—counteract the greater connectivity of females between sites.

Large-scale studies of song culture in the wild are challenging. In contrast to a laboratory situation, it is difficult to amass data, the identity of individuals is not known, and there is little control over the social context of singing—which limits assessment of individual-level mechanisms. However, our population-level analyses reveal distinct and complex female culture with different spatial patterns to male culture. This underlines the inadequacy of the male-centric research paradigm and calls for comparisons of male and female repertoires in many more species (Riebel et al., 2019). Sophisticated field experiments are now needed to resolve the mechanisms of dispersal, selection and learning modes that give rise to the pattern we have uncovered.

## Supporting information

Article S1

## Author contributions

Dianne Brunton and Wesley Webb conceived of the study. Wesley Webb, Michelle Roper, Dianne Brunton, and Aaron Harmer conducted the fieldwork with help from volunteers. Wesley Webb, Yukio Fukuzawa, and Michelle Roper created the song database. Wesley Webb, Yukio Fukuzawa and Matthew Pawley analysed the data, with guidance from Dianne Brunton and Aaron Harmer. Wesley Webb wrote the paper with guidance from the other authors.

## Data availability

The interactive *Koe* database can be accessed online at koe.io.ac.nz with username *korimako* and password Bellbird_Culture. See Data S3 for a table of all acoustic units and annotations, and Data S4 for a table of all songs and their metadata. Measurements of all syllables can be accessed at 10.5281/zenodo.5072580.

## Acknowledgements

We thank Ngati Wai and Ngati Manuhiri iwi for their support, and Barbara Evans, Jessica Patino-Perez, and Mehrnaz Tavasoli for their help with recording song. The research was funded by a Marsden Fund grant (13-MAU-004) from the Royal Society of New Zealand Te Apārangi. The manuscript was improved by feedback from Rebecca Webb.

