## Supplementary material for "Sexually distinct song cultures in a songbird metapopulation": Article S1

### S1A—Classification methodology

WHW compared the 20,700 syllables visually and aurally to classify them into types based on just-noticeable differences. This process was made possible by *Koe* web-based bioacoustics software, which we designed for expedited syllable classification. A tutorial case study for large-scale classification in *Koe* is presented in Fukuzawa et al. (2020); this explains the process we followed.

There was one group of syllable types that was not suitable for the chosen classification method: pure-tone, flat-contour whistle syllables (hereafter *pipe*s). Unlike other bellbird syllable types, *pipes* exhibit a wide range of fundamental frequencies and durations, giving an overall impression of continual gradation rather than discrete categories. WHW classified *pipes* based on the nearest note on the piano keyboard by aurally comparing playback of each *pipe* syllable in *Koe* to an online sine wave tone generator (http://www.szynalski.com/tone-generator/). Duration was ignored in the classification of *pipe* syllables.

### S1B—Extrapolating syllable diversity estimates for each site

The raw number of types recorded at each site are not directly comparable, due to the inevitable confounding of different sampling effort. To account for this, we used the statistical software *EstimateS* (Colwell, 2013) to produce syllable type accumulation curves and extrapolate the true number of syllable types at each site (*S*).

To do so, unit table data were exported from *Koe*. The data were arranged in an *Excel* Pivot Table to show counts of each syllable type, with syllable types as 'species' (columns) and songs as ‘samples’ (rows). In *EstimateS* we chose “sample-based incidence or abundance data”, with “one set of replicated sampling units (classic EstimateS input)”—samples as rows, with ‘species’ (syllable types) as columns. Under “sample order randomization for estimators and indices” we used the default of 100 runs (the number of randomizations). Under “Extrapolation of rarefaction curves” we chose to extrapolate to a total of 1000 samples (songs). Under “Estimation points (knots) for rarefaction and extrapolation” we chose to estimate at every sample. Other settings were left at their default values. Rarefaction curve graphs were then produced in *Excel* from the *S(est)* values column, with *S(est) 95% CI Lower Bound* and *S(est) 95% CI Upper Bound* columns providing the 95% confidence interval for the estimate, with the *x* axis displaying values for the first 500 songs.

### S1C—Testing for sex differences in degree of site–site repertoire overlap

To test for an overall sex difference in degree of site–site repertoire overlap, we used a two-tailed Sign Test (Dixon and Mood, 1946) as a non-parametric alternative to a paired t-test. To do this, for each pair of sites we compared the degree of male–male repertoire overlap with the degree of female–female repertoire overlap. If the male–male overlap was greater, the comparison was marked as ‘+’. If the female–female overlap was greater, the comparison was marked as ‘–’. If overall there is no difference in the degree of site-site repertoire overlap between sexes (the null hypothesis), then the counts of positive and negative signs are expected to be equal. If males tend to have greater overlap, there will be significantly more positive signs than negative, whereas if females tend to have greater overlap, there will be significantly more negative signs than positive.
