## Supplementary material for "Sexually distinct song cultures in a songbird metapopulation": Table S1

**Catalogue of all syllable classes from across the Hauraki Gulf metapopulation.**

(***Below***) Each row of the table represents one fine-scale syllable class. The broad-scale 'family' to which the class belongs is given in the leftmost column. Classes with an asterisk (*) indicate classes that are sung by both sexes. Counts of the fine-scale class at each site are given in the right-hand columns. For robustness, the fine-scale classes displayed have at least three occurrences within the sex. Site abbreviations are as follows: HAU, Hauturu; TAW, Tawharanui; TMI, Tiri; LAI, Lady Alice; REP, Repanga; PKI, Poor Knights. The spectrogram associated with each class in the table was hand-chosen in *Koe* based on clarity and class representativeness. Time and frequency scalebars are provided at the end of the male and female sub-tables.

### Catalogue of male syllable classes

| Broad-scale syllable class | Fine-scale syllable class | Spectrogram | HAU | TAW | TMI | LAI | REP | PKI | Total |
|---|---|---|---|---|---|---|---|---|---|
| Alarmy* | Alarmy(cheese2) | 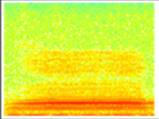 | 3 | 0 | 0 | 0 | 0 | 0 | **3** |
| Alarmy* | Alarmy(double) | 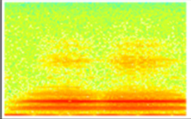 | 0 | 31 | 0 | 0 | 0 | 0 | **31** |
| Alarmy* | Alarmy(double+high) | 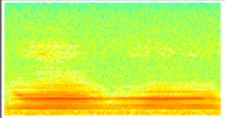 | 4 | 0 | 0 | 0 | 0 | 0 | **4** |
| Alarmy* | Alarmy(downflickstart) | 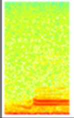 | 0 | 0 | 0 | 11 | 0 | 0 | **11** |
| Alarmy* | Alarmy(short) | 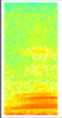 | 0 | 0 | 286 | 0 | 0 | 1 | **287** |
| Alarmy* | Alarmy(short+highbit)* | 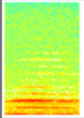 | 0 | 0 | 0 | 0 | 12 | 0 | **12** |
| Alarmy* | Alarmy(short+highbit1) | 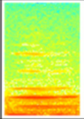 | 0 | 0 | 0 | 0 | 10 | 0 | **10** |
| Alarmy* | Alarmy(short+sharp) | 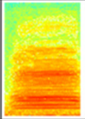 | 0 | 0 | 5 | 0 | 0 | 0 | **5** |

| Broad-scale syllable class | Fine-scale syllable class | Spectrogram | HAU | TAW | TMI | LAI | REP | PKI | Total |
|---|---|---|---|---|---|---|---|---|---|
| Alarmy* | Alarmy(short+up)* | 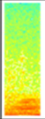 | 0 | 56 | 0 | 0 | 338 | 0 | **394** |
| Alarmy* | Alarmy(short+up+hah) | 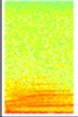 | 0 | 25 | 0 | 0 | 0 | 0 | **25** |
| Alarmy* | Alarmy(trainwhistle+aah) | 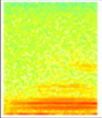 | 0 | 0 | 0 | 0 | 45 | 0 | **45** |
| Alarmy* | Alarmy(trainwhistle+ooh) | 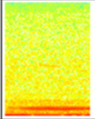 | 0 | 23 | 0 | 0 | 15 | 0 | **38** |
| Alarmy* | Alarmy(tritone1) | 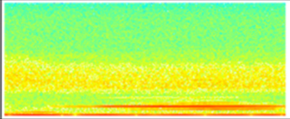 | 0 | 0 | 0 | 0 | 0 | 5 | **5** |
| Alarmy* | Alarmy(tritone2) | 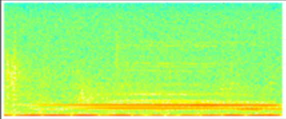 | 0 | 0 | 0 | 0 | 0 | 3 | **3** |
| Alarmy* | Alarmy(twobitCUV+beware)* | 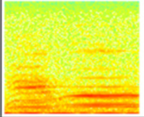 | 0 | 0 | 0 | 0 | 38 | 0 | **38** |
| Alarmy* | Alarmy(twobitCUV+hello) | 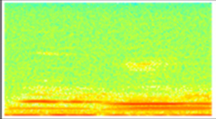 | 0 | 0 | 0 | 0 | 32 | 0 | **32** |
| Alarmy* | Alarmy(twobitCUV+io) | 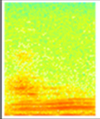 | 0 | 0 | 0 | 0 | 63 | 0 | **63** |

| Broad-scale syllable class | Fine-scale syllable class | Spectrogram | HAU | TAW | TMI | LAI | REP | PKI | Total |
|---|---|---|---|---|---|---|---|---|---|
| Alarmy* | Alarmy(twobitCUV+nihao) | 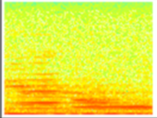 | 0 | 0 | 0 | 0 | 21 | 0 | **21** |
| Alarmy* | Alarmy(twobitCUV+rightio) | 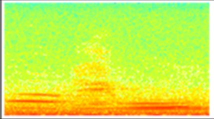 | 0 | 0 | 0 | 0 | 29 | 0 | **29** |
| Alarmy* | Alarmy(twobitCUV+stutter+beware) | 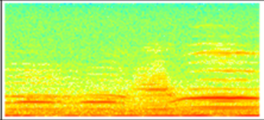 | 0 | 0 | 0 | 0 | 32 | 0 | **32** |
| Alarmy* | Alarmy(twobitCUV+stutter+rightio) | 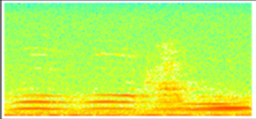 | 0 | 0 | 0 | 0 | 123 | 0 | **123** |
| Alarmy* | Alarmy_Pipe(high) | 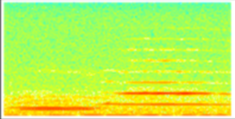 | 0 | 0 | 0 | 0 | 26 | 0 | **26** |
| Alarmy* | SimUpDown_Upsqueak(firsthalf) | 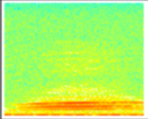 | 0 | 0 | 0 | 0 | 0 | 12 | **12** |
| Alarmy* | Stutter(alarmy)* | 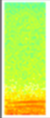 | 0 | 0 | 0 | 0 | 3 | 0 | **3** |
| Alarmy* | Upsqueak(alarmyLAI) | 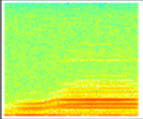 | 0 | 0 | 0 | 3 | 0 | 0 | **3** |
| Alarmy* | Upsqueak(kitten)* | 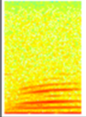 | 0 | 88 | 0 | 0 | 0 | 0 | **88** |

| Broad-scale syllable class | Fine-scale syllable class | Spectrogram | HAU | TAW | TMI | LAI | REP | PKI | Total |
|---|---|---|---|---|---|---|---|---|---|
| Alarmy* | Upsqueak(kitten)_Alarmy | 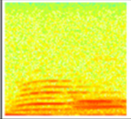 | 0 | 24 | 0 | 0 | 0 | 0 | **24** |
| Bip | Bip | 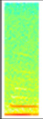 | 0 | 0 | 0 | 0 | 0 | 626 | **626** |
| Chiggle* | Chiggle* | 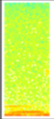 | 0 | 116 | 0 | 0 | 7 | 6 | **129** |
| Chiggle* | Chiggle(straight) | 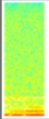 | 0 | 0 | 0 | 5 | 0 | 0 | **5** |
| Chortle* | Chortle(click) | 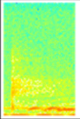 | 0 | 0 | 0 | 0 | 0 | 17 | **17** |
| Chortle* | Chortle(LAI) | 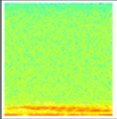 | 0 | 0 | 0 | 4 | 0 | 0 | **4** |
| Chortle* | Chortle(TAW) | 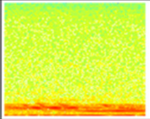 | 0 | 10 | 0 | 0 | 0 | 0 | **10** |
| Chortle* | Chortle(whipsong1) | 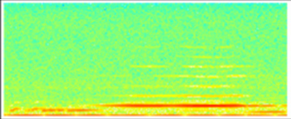 | 0 | 0 | 0 | 0 | 0 | 4 | **4** |
| Chortle* | Chortle(whipsong1+click) | 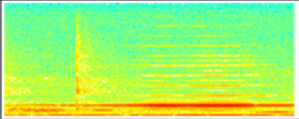 | 0 | 0 | 0 | 0 | 0 | 6 | **6** |

| Broad-scale syllable class | Fine-scale syllable class | Spectrogram | HAU | TAW | TMI | LAI | REP | PKI | Total |
|---|---|---|---|---|---|---|---|---|---|
| Chortle* | Chortle(whipsong2) | 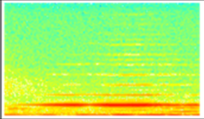 | 0 | 0 | 0 | 0 | 0 | 5 | **5** |
| Chortle* | Chortle(whipsong2+click) | 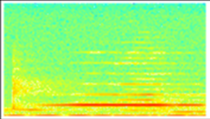 | 0 | 0 | 0 | 0 | 0 | 10 | **10** |
| Chortle* | Chortle_Ghh | 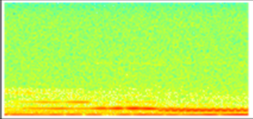 | 0 | 0 | 0 | 0 | 222 | 0 | **222** |
| Chortle* | Moustache(messy) | 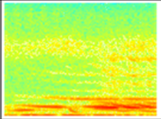 | 0 | 0 | 36 | 0 | 0 | 0 | **36** |
| Chortle* | Pipe(low)_Tailoff1 | 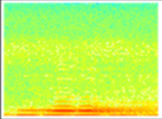 | 0 | 0 | 49 | 0 | 0 | 0 | **49** |
| Chortle* | Pipe(low)_Tailoff2 | 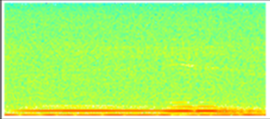 | 0 | 0 | 140 | 0 | 0 | 0 | **140** |
| Chump* | Chiup* | 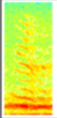 | 0 | 0 | 3 | 0 | 0 | 0 | **3** |
| Chump* | Chump | 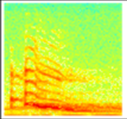 | 92 | 70 | 0 | 0 | 0 | 0 | **162** |
| Chump* | Chump(ratchety) | 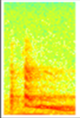 | 10 | 0 | 0 | 0 | 0 | 0 | **10** |

| Broad-scale syllable class | Fine-scale syllable class | Spectrogram | HAU | TAW | TMI | LAI | REP | PKI | Total |
|---|---|---|---|---|---|---|---|---|---|
| Chump* | Dragon | 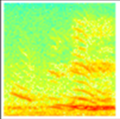 | 0 | 0 | 425 | 0 | 0 | 0 | **425** |
| Chump* | Ghh_Chump | 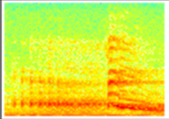 | 3 | 113 | 0 | 0 | 0 | 0 | **116** |
| Chump* | Ghh_Ghh_Chump | 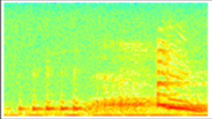 | 43 | 0 | 0 | 0 | 0 | 0 | **43** |
| Chump* | Handclasp(sloped) | 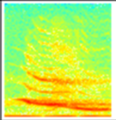 | 0 | 0 | 0 | 0 | 0 | 64 | **64** |
| Chump* | Pipe(low)_Bentpipe(LAI2) | 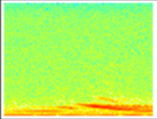 | 0 | 0 | 0 | 19 | 0 | 0 | **19** |
| Chump* | Pipe(rough)_Chump | 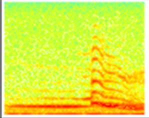 | 0 | 8 | 0 | 0 | 0 | 0 | **8** |
| Chump* | Rattle_Ghh | 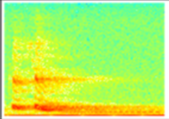 | 0 | 0 | 0 | 12 | 0 | 0 | **12** |
| Click | Click_Click(tail) | 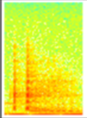 | 0 | 36 | 0 | 0 | 0 | 0 | **36** |
| Click | Pipe(low+click)_Tailoff | 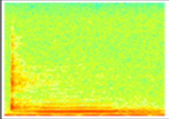 | 0 | 0 | 4 | 0 | 0 | 0 | **4** |

| Broad-scale syllable class | Fine-scale syllable class | Spectrogram | HAU | TAW | TMI | LAI | REP | PKI | Total |
|---|---|---|---|---|---|---|---|---|---|
| Cough* | Cough(clickCUV) | 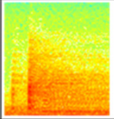 | 0 | 0 | 0 | 0 | 304 | 0 | **304** |
| Cough* | Cough(clickhigh)* | 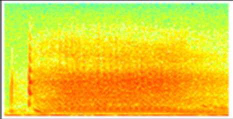 | 4 | 69 | 0 | 0 | 0 | 0 | **73** |
| Cough* | Cough(clicksquare) | 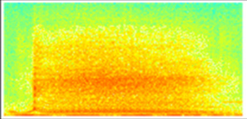 | 0 | 0 | 0 | 0 | 0 | 69 | **69** |
| Cough* | Cough(growl) | 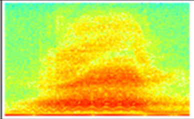 | 0 | 0 | 0 | 0 | 0 | 52 | **52** |
| Cough* | Cough(hiss) | 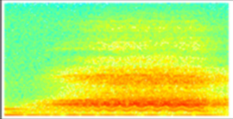 | 0 | 0 | 0 | 0 | 0 | 79 | **79** |
| Cough* | Cough(hiss+lower) | 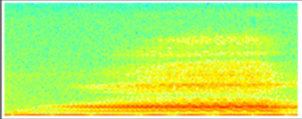 | 0 | 0 | 0 | 0 | 0 | 7 | **7** |
| Cough* | Cough(LAI) | 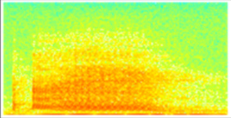 | 0 | 0 | 0 | 42 | 0 | 0 | **42** |
| Cough* | Cough(short) | 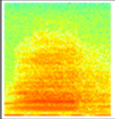 | 48 | 103 | 0 | 0 | 237 | 0 | **388** |
| Cough* | Cough(sneeze) | 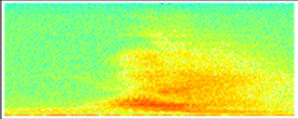 | 0 | 0 | 0 | 0 | 0 | 53 | **53** |

| Broad-scale syllable class | Fine-scale syllable class | Spectrogram | HAU | TAW | TMI | LAI | REP | PKI | Total |
|---|---|---|---|---|---|---|---|---|---|
| Cough* | Cough(TMI+click) | 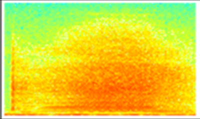 | 0 | 0 | 639 | 0 | 0 | 0 | **639** |
| Down* | Down(LAI1) | 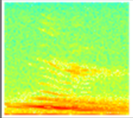 | 0 | 0 | 0 | 6 | 0 | 0 | **6** |
| Down* | Down(LAI2)* | 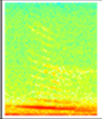 | 0 | 0 | 0 | 6 | 0 | 0 | **6** |
| Down* | Down(LAI3)* | 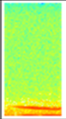 | 0 | 0 | 0 | 7 | 0 | 0 | **7** |
| Down* | Down(LBI1)* | 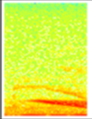 | 3 | 0 | 0 | 0 | 0 | 0 | **3** |
| Down* | Down(slightcurve+click)* | 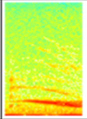 | 4 | 1 | 0 | 0 | 0 | 0 | **5** |
| Down* | Down(slightcurve1)* | 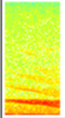 | 10 | 0 | 0 | 0 | 0 | 0 | **10** |
| Down* | Down_Tink | 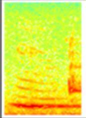 | 0 | 13 | 0 | 0 | 0 | 0 | **13** |
| Down* | Downsweep(roughtosmooth)* | 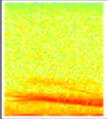 | 0 | 0 | 0 | 7 | 0 | 0 | **7** |

| Broad-scale syllable class | Fine-scale syllable class | Spectrogram | HAU | TAW | TMI | LAI | REP | PKI | Total |
|---|---|---|---|---|---|---|---|---|---|
| Down_Pipe* | Cascade | 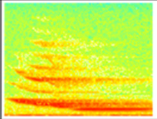 | 1 | 0 | 0 | 0 | 0 | 12 | **13** |
| Down_Pipe* | Down_Pipe(G#[6])* | 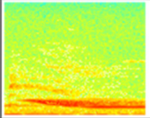 | 7 | 0 | 0 | 0 | 0 | 0 | **7** |
| Down_Pipe* | Down_Pipe(G[6]+)* | 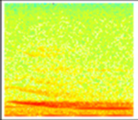 | 6 | 0 | 0 | 0 | 0 | 0 | **6** |
| Downsqueak* | Doubledownsqueak(LAI) | 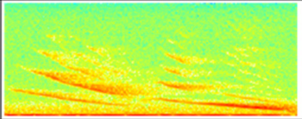 | 0 | 0 | 0 | 6 | 0 | 0 | **6** |
| Downsqueak* | Downsqueak(alarmy)_Tink | 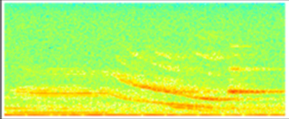 | 3 | 0 | 0 | 0 | 0 | 0 | **3** |
| Downsqueak* | Downsqueak(hook)* | 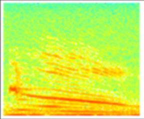 | 0 | 0 | 483 | 0 | 0 | 0 | **483** |
| Downsqueak* | Downsqueak(laser)* | 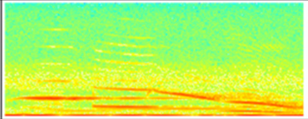 | 0 | 0 | 0 | 0 | 0 | 5 | **5** |
| Downsqueak* | Downsqueak_Downlines(LAI) | 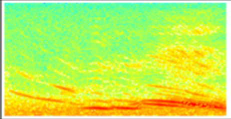 | 0 | 0 | 0 | 4 | 0 | 0 | **4** |
| Downsqueak* | Flatsqueak_Downsqueak_Tink | 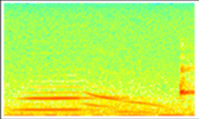 | 0 | 17 | 0 | 0 | 0 | 0 | **17** |

| Broad-scale syllable class | Fine-scale syllable class | Spectrogram | HAU | TAW | TMI | LAI | REP | PKI | Total |
|---|---|---|---|---|---|---|---|---|---|
| Downsqueak* | Flatsqueak_Downsqueak_Tinkee | 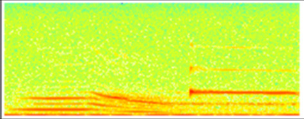 | 0 | 69 | 0 | 0 | 0 | 0 | **69** |
| Downsqueak* | Shrieky(uphook) | 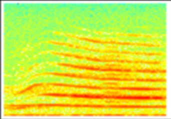 | 0 | 0 | 121 | 0 | 0 | 0 | **121** |
| Downsqueak* | Shrieky(uphook+lower) | 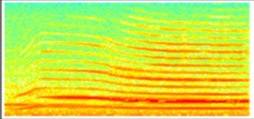 | 0 | 0 | 26 | 0 | 0 | 0 | **26** |
| Downsqueak* | Shrieky(uphook+sansuphook) | 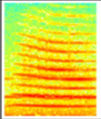 | 0 | 0 | 9 | 0 | 0 | 0 | **9** |
| Downsqueak_Pipe* | Downsqueak(bitunder)_Pipe(low) | 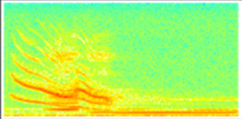 | 0 | 0 | 134 | 0 | 0 | 0 | **134** |
| Downsqueak_Pipe* | Downsqueak(bitunder)_Pipe(mid) | 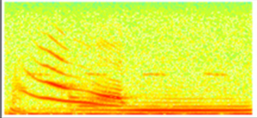 | 0 | 41 | 0 | 0 | 0 | 0 | **41** |
| Downsqueak_Pipe* | Downsqueak(chumpbit)_Pipe(high)* | 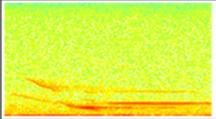 | 1 | 19 | 0 | 0 | 0 | 0 | **20** |
| Downsqueak_Pipe* | Downsqueak_BackwardsJ | 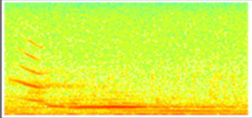 | 0 | 4 | 0 | 0 | 0 | 0 | **4** |
| Downsqueak_Pipe* | ysideways(D[6]) | 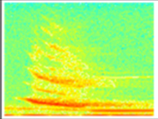 | 0 | 0 | 0 | 0 | 0 | 28 | **28** |

| Broad-scale syllable class | Fine-scale syllable class | Spectrogram | HAU | TAW | TMI | LAI | REP | PKI | Total |
|---|---|---|---|---|---|---|---|---|---|
| Downsqueak_Pipe* | ysideways(E[6]) | 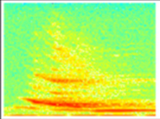 | 0 | 0 | 0 | 0 | 0 | 9 | **9** |
| Downsqueak_Pipe* | ysideways(F#[6]) | 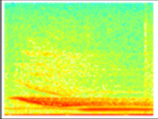 | 0 | 0 | 0 | 0 | 0 | 3 | **3** |
| Downsqueak_Pipe* | ysideways(G#[6]) | 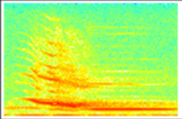 | 0 | 0 | 0 | 0 | 0 | 71 | **71** |
| Downsqueak_Stepdown | Teeyoo | 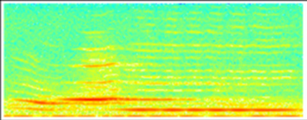 | 0 | 0 | 0 | 0 | 0 | 17 | **17** |
| Flatsqueak* | 8vepipe(A#[5]) | 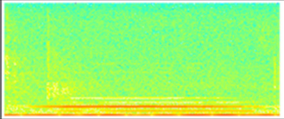 | 0 | 0 | 0 | 0 | 0 | 3 | **3** |
| Flatsqueak* | 8vepipe(A#[5]+) | 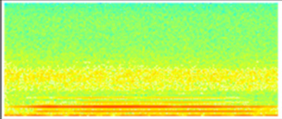 | 0 | 0 | 0 | 0 | 0 | 18 | **18** |
| Flatsqueak* | 8vepipe(B[5]) | 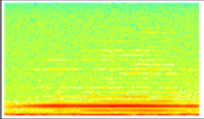 | 0 | 0 | 0 | 0 | 0 | 6 | **6** |
| Flatsqueak* | 8vepipe(F#[6]+) | 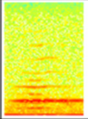 | 0 | 0 | 0 | 0 | 4 | 0 | **4** |
| Flatsqueak* | 8vepipe(F[6]) | 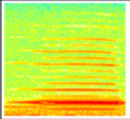 | 3 | 0 | 0 | 0 | 0 | 0 | **3** |

| Broad-scale syllable class | Fine-scale syllable class | Spectrogram | HAU | TAW | TMI | LAI | REP | PKI | Total |
|---|---|---|---|---|---|---|---|---|---|
| Flatsqueak* | Cheesy | 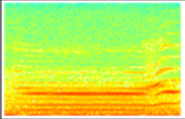 | 0 | 0 | 0 | 0 | 249 | 0 | **249** |
| Flatsqueak* | Down(jumpup)_Flatsqueak(1) | 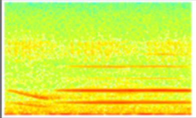 | 4 | 0 | 0 | 0 | 0 | 0 | **4** |
| Flatsqueak* | Downhook_Flatsqueak | 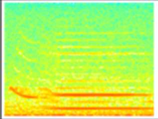 | 0 | 0 | 42 | 0 | 0 | 0 | **42** |
| Flatsqueak* | Downhook_Flatsqueak(short) | 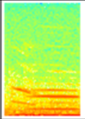 | 0 | 0 | 3 | 0 | 77 | 0 | **80** |
| Flatsqueak* | Flatsqueak(CUV+B) | 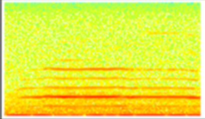 | 0 | 0 | 0 | 0 | 9 | 0 | **9** |
| Flatsqueak* | Flatsqueak(CUV+C#) | 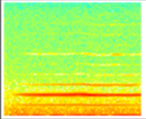 | 0 | 0 | 0 | 0 | 15 | 0 | **15** |
| Flatsqueak* | Flatsqueak(D[7]) | 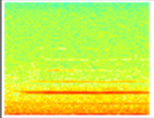 | 0 | 0 | 0 | 0 | 4 | 0 | **4** |
| Flatsqueak* | Flatsqueak(downhook) | 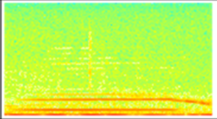 | 0 | 1 | 0 | 0 | 71 | 0 | **72** |
| Flatsqueak* | Flatsqueak(F#[6]) | 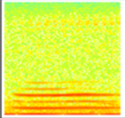 | 4 | 0 | 0 | 2 | 0 | 0 | **6** |

| Broad-scale syllable class | Fine-scale syllable class | Spectrogram | HAU | TAW | TMI | LAI | REP | PKI | Total |
|---|---|---|---|---|---|---|---|---|---|
| Flatsqueak* | Flatsqueak_Tink | 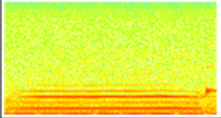 | 0 | 7 | 0 | 0 | 0 | 0 | **7** |
| Flatsqueak* | Flatsqueak_UFO | 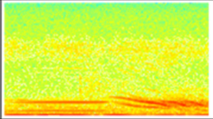 | 3 | 0 | 0 | 0 | 0 | 0 | **3** |
| Flatsqueak* | Upsqueak_Flatsqueak(2200Hz) | 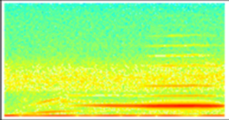 | 0 | 0 | 0 | 0 | 0 | 7 | **7** |
| Flatsqueak* | Upsqueak_Flatsqueak(3) | 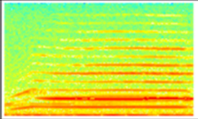 | 0 | 0 | 29 | 0 | 0 | 0 | **29** |
| Flatsqueak* | Upsqueak_Flatsqueak_Table | 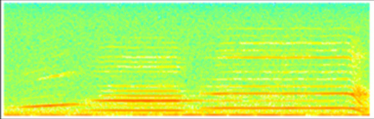 | 19 | 0 | 0 | 0 | 0 | 0 | **19** |
| Peaksqueak* | CUVsqueak* | 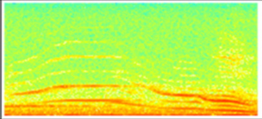 | 0 | 0 | 0 | 0 | 204 | 0 | **204** |
| Peaksqueak* | Down_Peak(LAI1) | 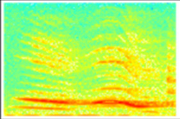 | 0 | 0 | 0 | 3 | 0 | 0 | **3** |
| Peaksqueak* | Kweeup | 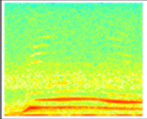 | 0 | 0 | 0 | 0 | 0 | 6 | **6** |
| Peaksqueak* | Peaksqueak(cutting)* | 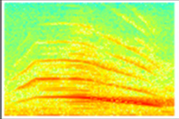 | 0 | 0 | 77 | 0 | 0 | 0 | **77** |

| Broad-scale syllable class | Fine-scale syllable class | Spectrogram | HAU | TAW | TMI | LAI | REP | PKI | Total |
|---|---|---|---|---|---|---|---|---|---|
| Peaksqueak* | Peaksqueak(whistley)* | 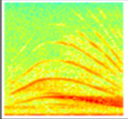 | 0 | 0 | 26 | 0 | 0 | 0 | **26** |
| Peaksqueak* | Pipe(low)_Weeyoo* | 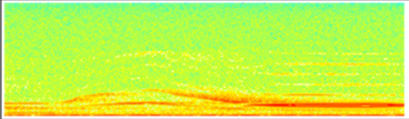 | 0 | 0 | 0 | 0 | 0 | 44 | **44** |
| Peaksqueak* | Woah* | 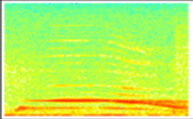 | 0 | 0 | 0 | 0 | 0 | 30 | **30** |
| Peaksqueak_Pipe | Peaksqueak_Pipe(E[5]) | 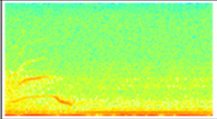 | 0 | 0 | 0 | 0 | 0 | 4 | **4** |
| Peaksqueak_Pipe | Wickoo | 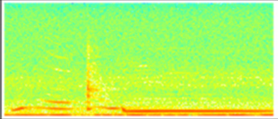 | 0 | 0 | 0 | 0 | 0 | 18 | **18** |
| Peaksqueak_Pipe | Wickoo_stepup | 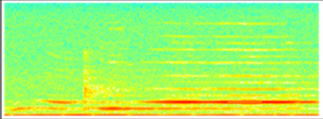 | 0 | 0 | 0 | 0 | 0 | 12 | **12** |
| Pipe* | Down(high)_Pipe(high) | 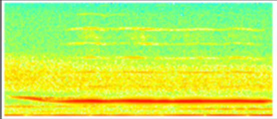 | 0 | 0 | 0 | 0 | 0 | 5 | **5** |
| Pipe* | Pipe(A#[5])* | 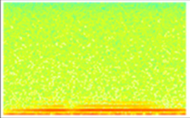 | 0 | 5 | 3 | 0 | 7 | 15 | **30** |
| Pipe* | Pipe(A#[5]+) | 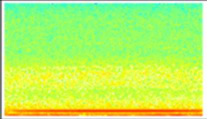 | 1 | 0 | 9 | 0 | 1 | 14 | **25** |

| Broad-scale syllable class | Fine-scale syllable class | Spectrogram | HAU | TAW | TMI | LAI | REP | PKI | Total |
|---|---|---|---|---|---|---|---|---|---|
| Pipe* | Pipe(A#[6])* | 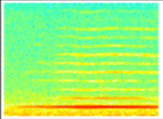 | 6 | 3 | 0 | 7 | 18 | 1 | **35** |
| Pipe* | Pipe(A#[6]+)* | 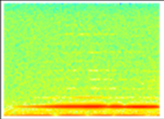 | 7 | 1 | 3 | 18 | 21 | 1 | **51** |
| Pipe* | Pipe(A#[6]+click)* | 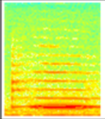 | 0 | 0 | 8 | 0 | 0 | 0 | **8** |
| Pipe* | Pipe(A#[6]bent)* | 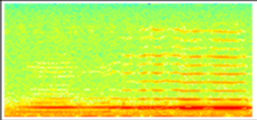 | 7 | 1 | 1 | 0 | 0 | 0 | **9** |
| Pipe* | Pipe(A[5]) | 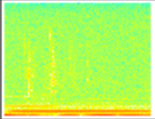 | 1 | 0 | 1 | 3 | 1 | 0 | **6** |
| Pipe* | Pipe(A[5]+) | 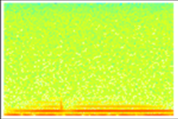 | 0 | 3 | 0 | 0 | 0 | 0 | **3** |
| Pipe* | Pipe(A[6])* | 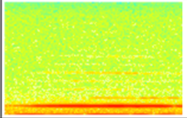 | 8 | 4 | 0 | 5 | 0 | 0 | **17** |
| Pipe* | Pipe(A[6]+)* | 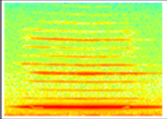 | 3 | 7 | 17 | 0 | 4 | 0 | **31** |
| Pipe* | Pipe(A[6]+bent) | 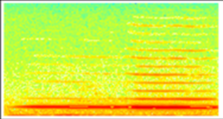 | 0 | 21 | 0 | 0 | 0 | 0 | **21** |

| Broad-scale syllable class | Fine-scale syllable class | Spectrogram | HAU | TAW | TMI | LAI | REP | PKI | Total |
|---|---|---|---|---|---|---|---|---|---|
| Pipe* | Pipe(A[6]+click) | 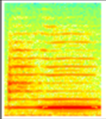 | 0 | 0 | 34 | 0 | 0 | 0 | **34** |
| Pipe* | Pipe(A[6]+click+) | 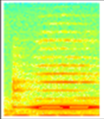 | 0 | 0 | 22 | 0 | 0 | 0 | **22** |
| Pipe* | Pipe(B[5])* | 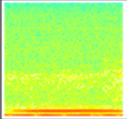 | 0 | 5 | 9 | 0 | 2 | 7 | **23** |
| Pipe* | Pipe(B[5]+) | 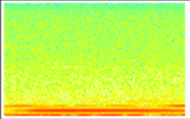 | 0 | 15 | 0 | 0 | 0 | 0 | **15** |
| Pipe* | Pipe(B[5]bentup) | 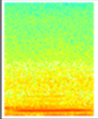 | 0 | 0 | 5 | 0 | 0 | 0 | **5** |
| Pipe* | Pipe(B[6])* | 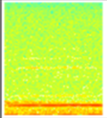 | 9 | 1 | 7 | 4 | 2 | 0 | **23** |
| Pipe* | Pipe(B[6]+)* | 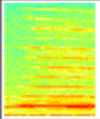 | 1 | 4 | 15 | 1 | 0 | 0 | **21** |
| Pipe* | Pipe(B[6]+click)* | 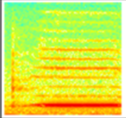 | 1 | 0 | 10 | 0 | 0 | 0 | **11** |
| Pipe* | Pipe(B[7])* | 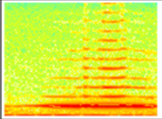 | 0 | 0 | 1 | 0 | 3 | 0 | **4** |

| Broad-scale syllable class | Fine-scale syllable class | Spectrogram | HAU | TAW | TMI | LAI | REP | PKI | Total |
|---|---|---|---|---|---|---|---|---|---|
| Pipe* | Pipe(C#[6])* | 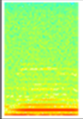 | 3 | 0 | 10 | 1 | 0 | 0 | **14** |
| Pipe* | Pipe(C#[6]+)* | 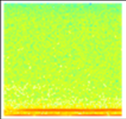 | 2 | 0 | 3 | 0 | 0 | 0 | **5** |
| Pipe* | Pipe(C#[6]bent) | 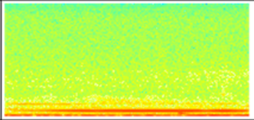 | 0 | 0 | 0 | 0 | 0 | 11 | **11** |
| Pipe* | Pipe(C#[7])* | 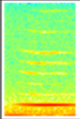 | 0 | 0 | 2 | 1 | 5 | 0 | **8** |
| Pipe* | Pipe(C#[7]+)* | 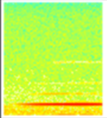 | 1 | 0 | 0 | 43 | 0 | 0 | **44** |
| Pipe* | Pipe(C#[7]+hook+click) | 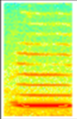 | 0 | 0 | 9 | 0 | 0 | 0 | **9** |
| Pipe* | Pipe(C[6]) | 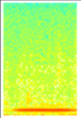 | 0 | 1 | 11 | 1 | 0 | 0 | **13** |
| Pipe* | Pipe(C[6]+) | 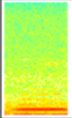 | 0 | 1 | 28 | 0 | 1 | 1 | **31** |
| Pipe* | Pipe(C[7])* | 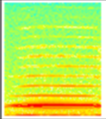 | 2 | 12 | 4 | 0 | 0 | 0 | **18** |

| Broad-scale syllable class | Fine-scale syllable class | Spectrogram | HAU | TAW | TMI | LAI | REP | PKI | Total |
|---|---|---|---|---|---|---|---|---|---|
| Pipe* | Pipe(C[7]+)* | 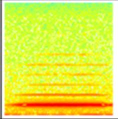 | 0 | 10 | 3 | 1 | 3 | 2 | **19** |
| Pipe* | Pipe(C[7]+click+) | 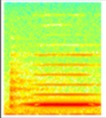 | 0 | 0 | 8 | 0 | 0 | 0 | **8** |
| Pipe* | Pipe(CUVbent) | 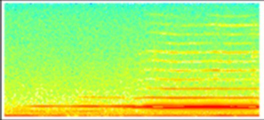 | 0 | 0 | 0 | 0 | 77 | 0 | **77** |
| Pipe* | Pipe(D#[6])* | 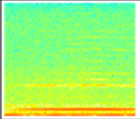 | 3 | 5 | 5 | 11 | 0 | 2 | **26** |
| Pipe* | Pipe(D#[6]+)* | 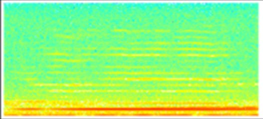 | 3 | 3 | 1 | 0 | 4 | 13 | **24** |
| Pipe* | Pipe(D#[6]bent2) | 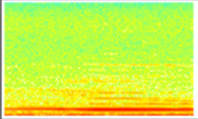 | 0 | 0 | 4 | 0 | 0 | 0 | **4** |
| Pipe* | Pipe(D#[7])* | 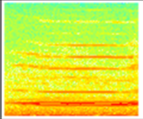 | 1 | 0 | 2 | 1 | 0 | 0 | **4** |
| Pipe* | Pipe(D#[7]+)* | 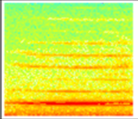 | 1 | 0 | 1 | 1 | 0 | 0 | **3** |
| Pipe* | Pipe(D[6])* | 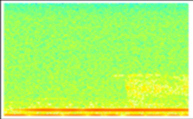 | 4 | 6 | 6 | 0 | 0 | 0 | **16** |

| Broad-scale syllable class | Fine-scale syllable class | Spectrogram | HAU | TAW | TMI | LAI | REP | PKI | Total |
|---|---|---|---|---|---|---|---|---|---|
| Pipe* | Pipe(D[6]+)* | 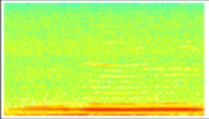 | 5 | 19 | 12 | 1 | 0 | 2 | **39** |
| Pipe* | Pipe(D[6]bent) | 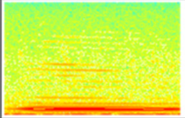 | 1 | 9 | 4 | 0 | 0 | 0 | **14** |
| Pipe* | Pipe(D[7])* | 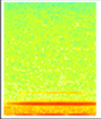 | 7 | 2 | 0 | 36 | 0 | 0 | **45** |
| Pipe* | Pipe(D[7]+)* | 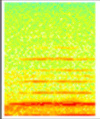 | 5 | 0 | 0 | 4 | 1 | 0 | **10** |
| Pipe* | Pipe(E[5]) | 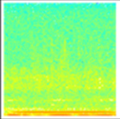 | 0 | 0 | 0 | 0 | 0 | 5 | **5** |
| Pipe* | Pipe(E[6])* | 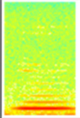 | 3 | 2 | 4 | 3 | 32 | 28 | **72** |
| Pipe* | Pipe(E[6]+)* | 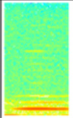 | 1 | 0 | 10 | 0 | 175 | 14 | **200** |
| Pipe* | Pipe(E[6]bent) | 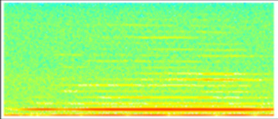 | 3 | 0 | 0 | 0 | 5 | 19 | **27** |
| Pipe* | Pipe(E[6]bent+click) | 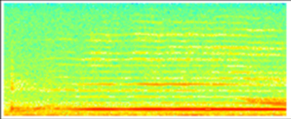 | 0 | 0 | 0 | 0 | 0 | 22 | **22** |

| Broad-scale syllable class | Fine-scale syllable class | Spectrogram | HAU | TAW | TMI | LAI | REP | PKI | Total |
|---|---|---|---|---|---|---|---|---|---|
| Pipe* | Pipe(E[7]bent)* | 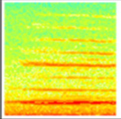 | 0 | 0 | 2 | 0 | 0 | 1 | **3** |
| Pipe* | Pipe(F#[5]) | 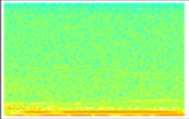 | 4 | 0 | 0 | 0 | 0 | 16 | **20** |
| Pipe* | Pipe(F#[6])* | 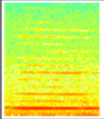 | 5 | 3 | 12 | 2 | 26 | 1 | **49** |
| Pipe* | Pipe(F#[6]+)* | 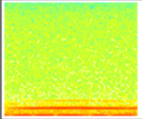 | 4 | 40 | 26 | 2 | 24 | 2 | **98** |
| Pipe* | Pipe(F#[6]bent) | 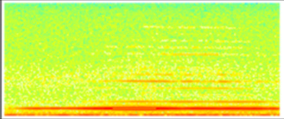 | 0 | 0 | 0 | 1 | 0 | 2 | **3** |
| Pipe* | Pipe(F#[6]bent+click) | 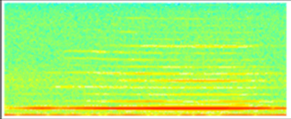 | 0 | 0 | 0 | 0 | 0 | 16 | **16** |
| Pipe* | Pipe(F#[7])* | 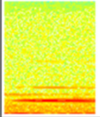 | 4 | 0 | 0 | 1 | 1 | 1 | **7** |
| Pipe* | Pipe(F#[7]+)* | 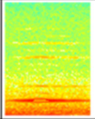 | 1 | 1 | 0 | 0 | 5 | 0 | **7** |
| Pipe* | Pipe(F[5]) | 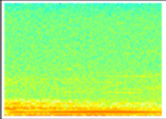 | 2 | 0 | 1 | 0 | 0 | 32 | **35** |

| Broad-scale syllable class | Fine-scale syllable class | Spectrogram | HAU | TAW | TMI | LAI | REP | PKI | Total |
|---|---|---|---|---|---|---|---|---|---|
| Pipe* | Pipe(F[5]+) | 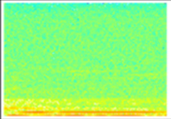 | 0 | 0 | 0 | 0 | 0 | 10 | **10** |
| Pipe* | Pipe(F[6])* | 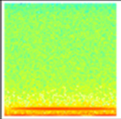 | 4 | 0 | 2 | 2 | 133 | 6 | **147** |
| Pipe* | Pipe(F[6]+)* | 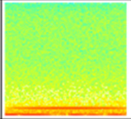 | 1 | 0 | 1 | 1 | 24 | 0 | **27** |
| Pipe* | Pipe(F[6]+click) | 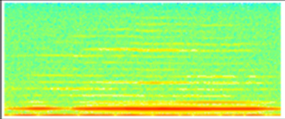 | 0 | 0 | 0 | 0 | 0 | 44 | **44** |
| Pipe* | Pipe(F[6]+click+) | 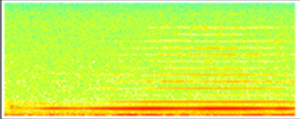 | 0 | 0 | 0 | 0 | 0 | 13 | **13** |
| Pipe* | Pipe(F[6]+double+) | 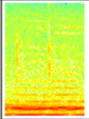 | 0 | 4 | 0 | 0 | 0 | 0 | **4** |
| Pipe* | Pipe(F[6]bent) | 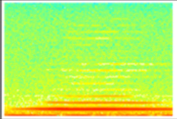 | 0 | 0 | 6 | 0 | 35 | 8 | **49** |
| Pipe* | Pipe(F[7]+)* | 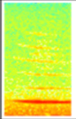 | 0 | 0 | 0 | 0 | 4 | 1 | **5** |
| Pipe* | Pipe(F[7]bent)* | 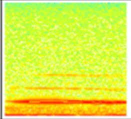 | 3 | 0 | 1 | 0 | 0 | 0 | **4** |

| Broad-scale syllable class | Fine-scale syllable class | Spectrogram | HAU | TAW | TMI | LAI | REP | PKI | Total |
|---|---|---|---|---|---|---|---|---|---|
| Pipe* | Pipe(G#[5]) | 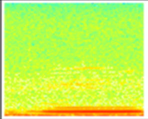 | 0 | 0 | 15 | 1 | 0 | 0 | **16** |
| Pipe* | Pipe(G#[6])* | 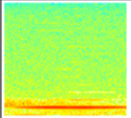 | 0 | 4 | 3 | 2 | 1 | 0 | **10** |
| Pipe* | Pipe(G#[6]+)* | 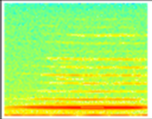 | 8 | 15 | 1 | 2 | 0 | 1 | **27** |
| Pipe* | Pipe(G#[6]+click) | 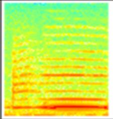 | 0 | 0 | 3 | 0 | 0 | 0 | **3** |
| Pipe* | Pipe(G#[6]+click+) | 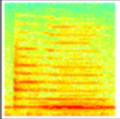 | 0 | 0 | 51 | 0 | 0 | 0 | **51** |
| Pipe* | Pipe(G#[6]+double) | 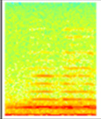 | 0 | 7 | 0 | 0 | 0 | 0 | **7** |
| Pipe* | Pipe(G#[6]+hard) | 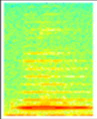 | 0 | 0 | 4 | 0 | 0 | 0 | **4** |
| Pipe* | Pipe(G#[6]bent)* | 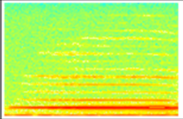 | 1 | 0 | 0 | 2 | 0 | 5 | **8** |
| Pipe* | Pipe(G#[6]rough) | 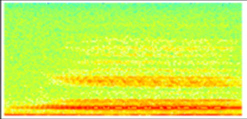 | 0 | 0 | 0 | 0 | 0 | 12 | **12** |

| Broad-scale syllable class | Fine-scale syllable class | Spectrogram | HAU | TAW | TMI | LAI | REP | PKI | Total |
|---|---|---|---|---|---|---|---|---|---|
| Pipe* | Pipe(G[5]+) | 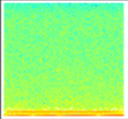 | 0 | 0 | 1 | 2 | 0 | 1 | **4** |
| Pipe* | Pipe(G[5]+click) | 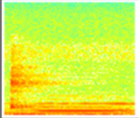 | 0 | 0 | 7 | 0 | 0 | 0 | **7** |
| Pipe* | Pipe(G[5]+click+) | 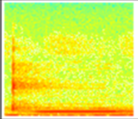 | 0 | 0 | 7 | 0 | 0 | 0 | **7** |
| Pipe* | Pipe(G[6])* | 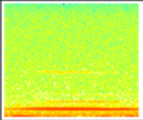 | 3 | 26 | 15 | 8 | 0 | 6 | **58** |
| Pipe* | Pipe(G[6]+)* | 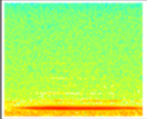 | 3 | 5 | 9 | 12 | 1 | 2 | **32** |
| Pipe* | Pipe(G[6]+rough) | 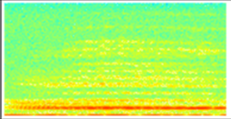 | 0 | 0 | 0 | 0 | 0 | 18 | **18** |
| Pipe* | Pipe(G[6]bent) | 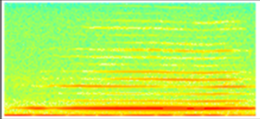 | 0 | 5 | 0 | 8 | 0 | 45 | **58** |
| Pipe* | Pipe(G[6]rough) | 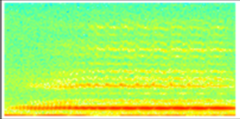 | 0 | 0 | 0 | 0 | 0 | 17 | **17** |
| Pipe* | Pipe(G[6]rough+) | 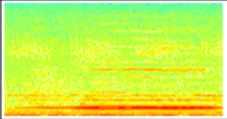 | 0 | 0 | 3 | 0 | 0 | 0 | **3** |

| Broad-scale syllable class | Fine-scale syllable class | Spectrogram | HAU | TAW | TMI | LAI | REP | PKI | Total |
|---|---|---|---|---|---|---|---|---|---|
| Pipe* | Pipe(hard)_Tink(A[6]) | 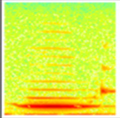 | 0 | 29 | 0 | 0 | 0 | 0 | **29** |
| Pipe* | Pipe(hard)_Tink(G#[6]+) | 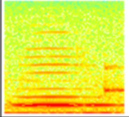 | 0 | 31 | 0 | 0 | 0 | 0 | **31** |
| Pipe* | Pipe(hard)_Tink(G[6]+) | 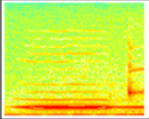 | 0 | 4 | 0 | 0 | 0 | 0 | **4** |
| Pipe* | Pipe(mid+tinystepup2) | 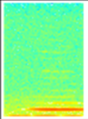 | 0 | 0 | 0 | 0 | 5 | 0 | **5** |
| Pipe* | Pipe(roughCUV1) | 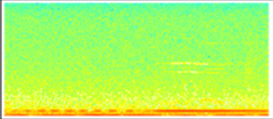 | 0 | 0 | 0 | 0 | 75 | 0 | **75** |
| Pipe* | Pipe(roughCUV2) | 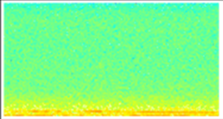 | 0 | 0 | 0 | 0 | 5 | 0 | **5** |
| Pipe* | Pipe(roughCUV3) | 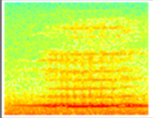 | 0 | 0 | 0 | 0 | 4 | 0 | **4** |
| Pipe* | Pipe(trilledLAI2) | 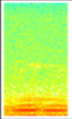 | 0 | 0 | 0 | 3 | 0 | 0 | **3** |
| Pipe* | Pipe(trilledLAI3) | 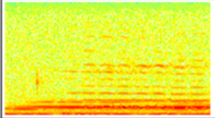 | 0 | 0 | 0 | 17 | 0 | 0 | **17** |

| Broad-scale syllable class | Fine-scale syllable class | Spectrogram | HAU | TAW | TMI | LAI | REP | PKI | Total |
|---|---|---|---|---|---|---|---|---|---|
| Pipe* | Wickee | 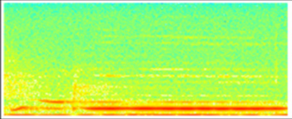 | 0 | 0 | 0 | 0 | 0 | 4 | **4** |
| Pipe_Cough | Pipe(A#[5])_Cough | 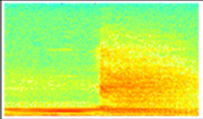 | 0 | 0 | 0 | 0 | 0 | 22 | **22** |
| Pipe_Cough | Pipe(B[5])_Cough | 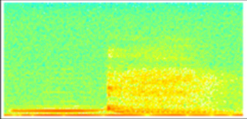 | 0 | 0 | 0 | 0 | 0 | 95 | **95** |
| Pipe_Cough | Pipe(B[5]+)_Cough | 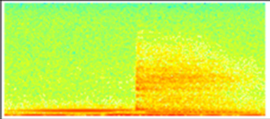 | 0 | 0 | 0 | 0 | 0 | 69 | **69** |
| Pipe_Cough | Pipe(rough+F#[6])_Cough | 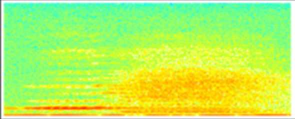 | 0 | 0 | 0 | 0 | 0 | **4** | **4** |
| Pipe_Cough | Pipe(rough+G#[6])_Cough | 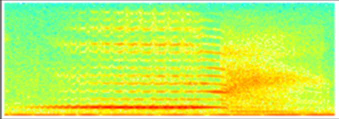 | 0 | 0 | 0 | 0 | 0 | 5 | **5** |
| Pipe_Cough | Pipe(rough+G#[6]+)_Cough | 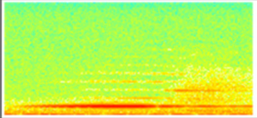 | 0 | 0 | 0 | 0 | 0 | 3 | **3** |
| Pipe_Cough | Pipe(rough+G[6])_Cough | 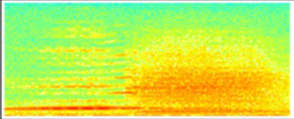 | 0 | 0 | 0 | 0 | 0 | 63 | **63** |
| Pipe_Cough | Pipe(rough+G[6]+)_Cough | 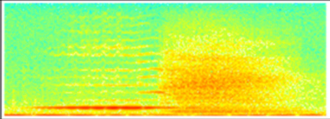 | 0 | 0 | 0 | 0 | 0 | 6 | **6** |

| Broad-scale syllable class | Fine-scale syllable class | Spectrogram | HAU | TAW | TMI | LAI | REP | PKI | Total |
|---|---|---|---|---|---|---|---|---|---|
| Pipe_Down* | Handclasp(flat) | 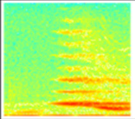 | 0 | 0 | 0 | 0 | 0 | 40 | **40** |
| Pipe_Down* | Pipe(E[7])_Dropoff* | 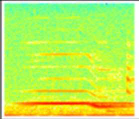 | 0 | 0 | 0 | 4 | 0 | 1 | **5** |
| Pipe_Down* | Pipe_Down_Tink | 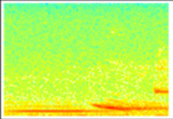 | 11 | 0 | 0 | 0 | 0 | 0 | **11** |
| Pipe_Downsqueak* | Pipe_Downlines | 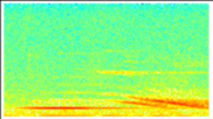 | 0 | 0 | 0 | 29 | 0 | 0 | **29** |
| Pipe_Downsqueak* | Pipe_Downlines(overlapping)_Tink | 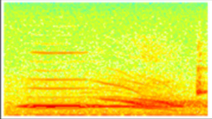 | 0 | 15 | 0 | 0 | 0 | 0 | **15** |
| Pipe_Downsqueak* | Pipe_Downsqueak* | 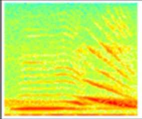 | 0 | 0 | 42 | 0 | 0 | 0 | **42** |
| Pipe_Downsqueak* | Pipe_Downsqueak_Jumpup | 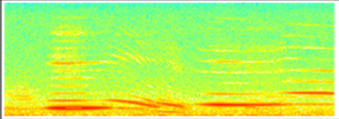 | 0 | 0 | 15 | 0 | 0 | 0 | **15** |
| Pipe_Downsqueak* | Pipe_Downsqueak2 | 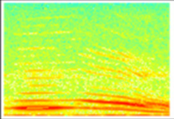 | 0 | 0 | 29 | 0 | 0 | 0 | **29** |
| Pipe_Downsqueak* | Pipe_Downsqueak4 | 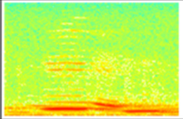 | 0 | 0 | 3 | 0 | 0 | 0 | **3** |

| Broad-scale syllable class | Fine-scale syllable class | Spectrogram | HAU | TAW | TMI | LAI | REP | PKI | Total |
|---|---|---|---|---|---|---|---|---|---|
| RoughBuzz* | Buzz(low) | 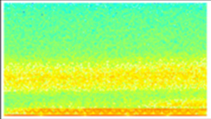 | 0 | 0 | 0 | 0 | 0 | 36 | **36** |
| RoughBuzz* | Buzz(shriek) | 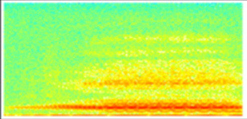 | 0 | 0 | 0 | 0 | 0 | 4 | **4** |
| RoughBuzz* | Crescendo(rough)* | 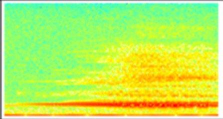 | 0 | 0 | 0 | 19 | 0 | 0 | **19** |
| RoughBuzz* | PKIShriek | 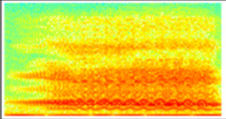 | 0 | 0 | 0 | 0 | 0 | 12 | **12** |
| RoughBuzz* | Spiderfangs | 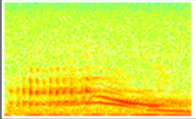 | 4 | 0 | 0 | 0 | 0 | 0 | **4** |
| Shrieky* | Upsqueak(shrieky)2 | 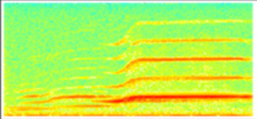 | 5 | 0 | 0 | 0 | 0 | 0 | **5** |
| Shrieky* | Upsqueak(shrieky+lower) | 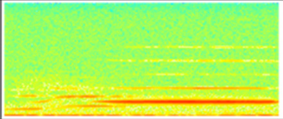 | 4 | 0 | 0 | 0 | 0 | 0 | **4** |
| Stepdown* | Burger(LBI) | 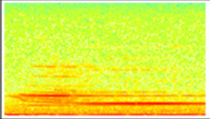 | 3 | 0 | 0 | 0 | 0 | 0 | **3** |
| Stepdown* | DinnerplateFlatArm* | 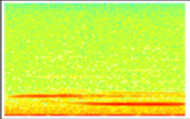 | 6 | 0 | 0 | 0 | 0 | 0 | **6** |

| Broad-scale syllable class | Fine-scale syllable class | Spectrogram | HAU | TAW | TMI | LAI | REP | PKI | Total |
|---|---|---|---|---|---|---|---|---|---|
| Stepdown* | Down(jumpup)_Stepdown | 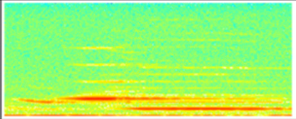 | 0 | 0 | 0 | 0 | 0 | 3 | **3** |
| Stepdown* | Pipe(mid+tinystepdown1) | 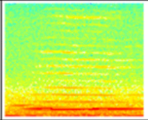 | 0 | 0 | 0 | 0 | 6 | 0 | **6** |
| Stepdown* | Stepdown(A#[6]-G[5]+click) | 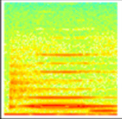 | 0 | 0 | 4 | 0 | 0 | 0 | **4** |
| Stepdown* | Stepdown(C#[7]-A[6]+click) | 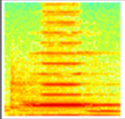 | 0 | 0 | 3 | 0 | 0 | 0 | **3** |
| Stepdown* | Stepdown(C[8]-D#[7]) | 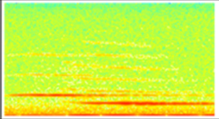 | 3 | 0 | 0 | 0 | 0 | 0 | **3** |
| Stepdown* | Stepdown(F#[7]-D#[6]) | 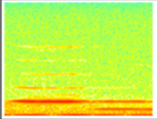 | 4 | 0 | 0 | 0 | 0 | 0 | **4** |
| Stepdown* | Stepdown(G#[6]-B[5]) | 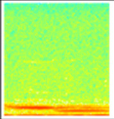 | 0 | 0 | 0 | 5 | 0 | 0 | **5** |
| Stepdown* | Stepdown(G[7]-D#[6])* | 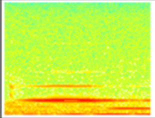 | 3 | 0 | 0 | 0 | 0 | 0 | **3** |
| Stepup* | Pipe(double)_8vepipe | 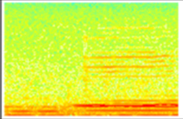 | 0 | 12 | 0 | 0 | 0 | 0 | **12** |

| Broad-scale syllable class | Fine-scale syllable class | Spectrogram | HAU | TAW | TMI | LAI | REP | PKI | Total |
|---|---|---|---|---|---|---|---|---|---|
| Stepup* | Pipe(trilled+LAI)_Pipe(A#[7]) | 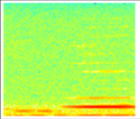 | 0 | 0 | 0 | 4 | 0 | 0 | **4** |
| Stepup* | Stepup(A#[5]-A[6]) | 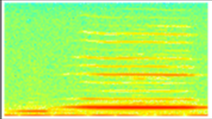 | 3 | 0 | 0 | 0 | 0 | 0 | **3** |
| Stepup* | Stepup(A[5]double-B[6]) | 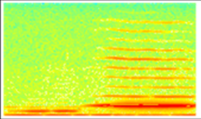 | 0 | 14 | 0 | 0 | 0 | 0 | **14** |
| Stepup* | Stepup(A[5]double-B[6]+) | 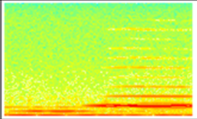 | 0 | 4 | 0 | 0 | 0 | 0 | **4** |
| Stepup* | Stepup(C[6]-C[7]) | 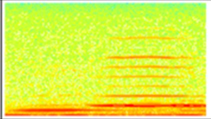 | 0 | 3 | 0 | 0 | 0 | 0 | **3** |
| Stepup* | Stepup(C[7]-D[7]) | 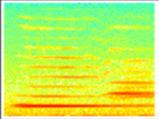 | 0 | 0 | 6 | 0 | 0 | 0 | **6** |
| Stepup* | Stepup(D[7]-D#[7]) | 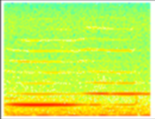 | 0 | 0 | 26 | 0 | 0 | 0 | **26** |
| Stepup* | Stepup(G#[5]-A[6]) | 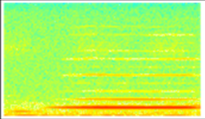 | 5 | 0 | 0 | 0 | 0 | 0 | **5** |
| Stepup_Down* | Pipe(low)_Bentpipe(LAI) | 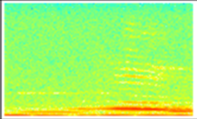 | 0 | 0 | 0 | 8 | 0 | 0 | **8** |

| Broad-scale syllable class | Fine-scale syllable class | Spectrogram | HAU | TAW | TMI | LAI | REP | PKI | Total |
|---|---|---|---|---|---|---|---|---|---|
| Stepup_Down* | Stepup_Slidedown | 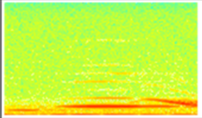 | 4 | 0 | 0 | 0 | 0 | 0 | **4** |
| Stutter* | Kweeup(short) | 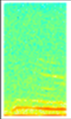 | 0 | 0 | 0 | 0 | 0 | 14 | **14** |
| Stutter* | Stutter* | 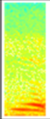 | 62 | 7 | 253 | 34 | 3 | 57 | **416** |
| Stutter* | Stutter(cheeselow) | 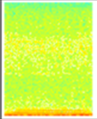 | 0 | 0 | 14 | 0 | 0 | 0 | **14** |
| Stutter* | Stutter(click)* | 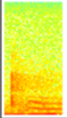 | 0 | 9 | 1 | 0 | 0 | 0 | **10** |
| Stutter* | Stutter(decurved)* | 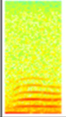 | 0 | 0 | 0 | 0 | 249 | 0 | **249** |
| Stutter* | Stutter(high)* | 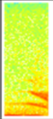 | 3 | 0 | 0 | 0 | 0 | 0 | **3** |
| Stutter* | Stutter(TAW)* | 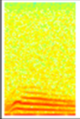 | 0 | 152 | 0 | 0 | 0 | 0 | **152** |
| Stutter(repeats)* | Stutter_Stutter(CUV+haha) | 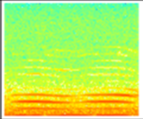 | 0 | 0 | 0 | 0 | 39 | 0 | **39** |

| Broad-scale syllable class | Fine-scale syllable class | Spectrogram | HAU | TAW | TMI | LAI | REP | PKI | Total |
|---|---|---|---|---|---|---|---|---|---|
| Stutter(repeats)* | Stutter_Stutter(LBITAW+nownow)* | 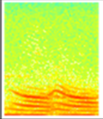 | 6 | 72 | 0 | 0 | 0 | 0 | **78** |
| Stutter(repeats)* | Stutter_Stutter(TMI+hoha) | 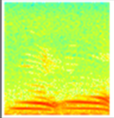 | 0 | 0 | 110 | 0 | 0 | 0 | **110** |
| Stutter(repeats)* | Stutter_Stutter_Stutter | 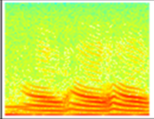 | 1 | 4 | 0 | 0 | 0 | 0 | **5** |
| Stutter_Pipe* | Down(carunlock) | 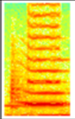 | 0 | 0 | 43 | 0 | 0 | 0 | **43** |
| Stutter_Stutter_Something | Stutter_Stutter_Pipe_Tink | 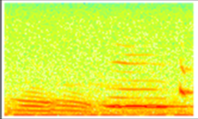 | 0 | 3 | 0 | 0 | 0 | 0 | **3** |
| Stutter_Stutter_Something | Stutter_Stutter_Upsqueak | 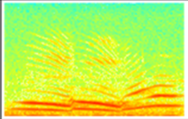 | 0 | 0 | 7 | 0 | 0 | 0 | **7** |
| Tink | Tink | 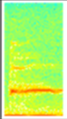 | 3 | 76 | 0 | 0 | 0 | 0 | **79** |
| Trill | Downdown(LAI) | 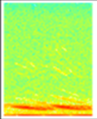 | 0 | 0 | 0 | 4 | 0 | 0 | **4** |
| Trill | Downlines(complex) | 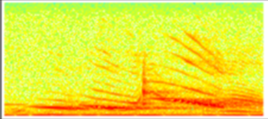 | 0 | 0 | 0 | 12 | 0 | 0 | **12** |

| Broad-scale syllable class | Fine-scale syllable class | Spectrogram | HAU | TAW | TMI | LAI | REP | PKI | Total |
|---|---|---|---|---|---|---|---|---|---|
| Trill | Downlines(complex2) | 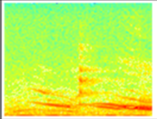 | 0 | 0 | 0 | 4 | 0 | 0 | **4** |
| Trill | Downlines(overlapping)_Tink | 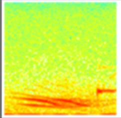 | 0 | 6 | 0 | 0 | 0 | 0 | **6** |
| Trill | Downlines(overlappingLAI1) | 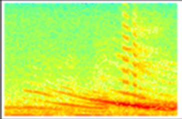 | 0 | 0 | 0 | 25 | 0 | 0 | **25** |
| Trill | Downlines(overlappingLAI2) | 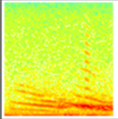 | 0 | 0 | 0 | 9 | 0 | 0 | **9** |
| Trill | Downlines(overlappingTAW1) | 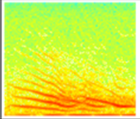 | 0 | 9 | 0 | 0 | 0 | 0 | **9** |
| Trill | Dragon(trill) | 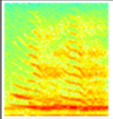 | 0 | 0 | 32 | 0 | 0 | 0 | **32** |
| Trill | Dragon(trill2) | 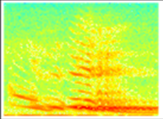 | 0 | 0 | 3 | 0 | 0 | 0 | **3** |
| Trill | Moustache(trill) | 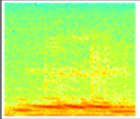 | 0 | 0 | 13 | 0 | 0 | 0 | **13** |
| Trill | Pipe(trilledLAI1) | 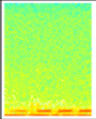 | 0 | 0 | 0 | 12 | 0 | 0 | **12** |

| Broad-scale syllable class | Fine-scale syllable class | Spectrogram | HAU | TAW | TMI | LAI | REP | PKI | Total |
|---|---|---|---|---|---|---|---|---|---|
| Trill | Rattle | 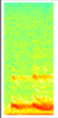 | 0 | 0 | 0 | 13 | 0 | 0 | **13** |
| Trill | Trill | 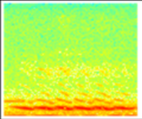 | 0 | 0 | 88 | 0 | 0 | 0 | **88** |
| Trill | Trill(rattle) | 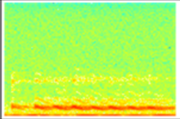 | 0 | 0 | 0 | 27 | 0 | 0 | **27** |
| Trill | Trill(short+notail) | 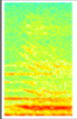 | 0 | 0 | 10 | 0 | 0 | 0 | **10** |
| Trill | Trill(short+twolines) | 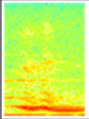 | 0 | 0 | 40 | 0 | 0 | 0 | **40** |
| Trill | Trill(tail) | 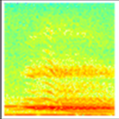 | 0 | 0 | 69 | 0 | 0 | 0 | **69** |
| Trill | Trill(tail+morebits) | 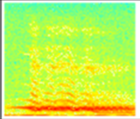 | 0 | 0 | 6 | 0 | 0 | 0 | **6** |
| Trill | UFO | 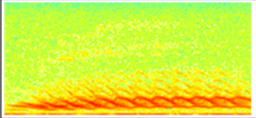 | 0 | 28 | 0 | 0 | 0 | 0 | **28** |
| Trill_Pipe* | Rattle_Pipe | 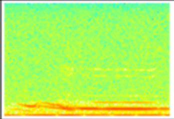 | 0 | 0 | 0 | 8 | 0 | 0 | **8** |

| Broad-scale syllable class | Fine-scale syllable class | Spectrogram | HAU | TAW | TMI | LAI | REP | PKI | Total |
|---|---|---|---|---|---|---|---|---|---|
| Trill_Pipe* | Snore | 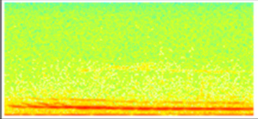 | 0 | 7 | 0 | 0 | 0 | 0 | **7** |
| Up | PKIWhip | 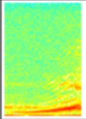 | 0 | 0 | 0 | 0 | 0 | 129 | **129** |
| Upsqueak* | SimUpDown_Upsqueak* | 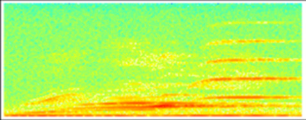 | 0 | 0 | 0 | 0 | 0 | 11 | **11** |
| Upsqueak* | Upsqueak(cleanglass)* | 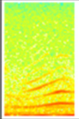 | 5 | 29 | 1 | 0 | 0 | 0 | **35** |
| Upsqueak* | Upsqueak(crazy+stepped) | 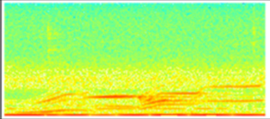 | 0 | 0 | 0 | 0 | 0 | 5 | **5** |
| Upsqueak* | Upsqueak(harmonic)* | 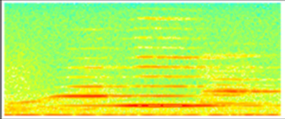 | 9 | 0 | 0 | 0 | 0 | 0 | **9** |
| Upsqueak* | Upsqueak(piercing) | 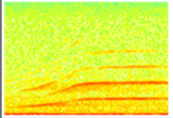 | 0 | 35 | 0 | 0 | 0 | 0 | **35** |
| Upsqueak* | Upsqueak(shrieky) | 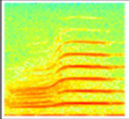 | 0 | 0 | 57 | 0 | 0 | 0 | **57** |
| Upsqueak_Flatsqueak | Upsqueak(stepped+suddenloud) | 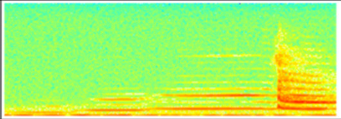 | 0 | 0 | 0 | 12 | 0 | 0 | **12** |

| Broad-scale syllable class | Fine-scale syllable class | Spectrogram | HAU | TAW | TMI | LAI | REP | PKI | Total |
|---|---|---|---|---|---|---|---|---|---|
| Upsqueak_Flatsqueak | Upsqueak(stepped+suddenloud2) | 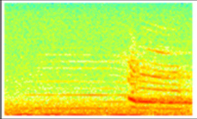 | 0 | 0 | 0 | 3 | 0 | 0 | **3** |
| Upsqueak_Flatsqueak | Upsqueak(twopeaks)_Pipe_Table | 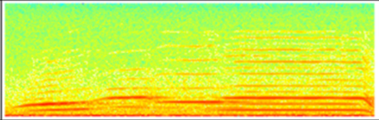 | 0 | 38 | 0 | 0 | 0 | 0 | **38** |
| Upsqueak_Flatsqueak | Upsqueak(twopeaks)_Pipe_Table(higher) | 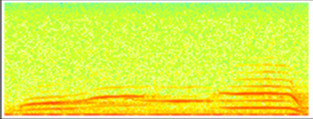 | 0 | 6 | 0 | 0 | 0 | 0 | **6** |
| Waah* | Cat(LAI) | 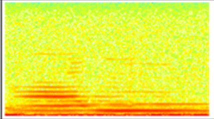 | 0 | 0 | 0 | 6 | 0 | 0 | **6** |
| Waah* | Kitten | 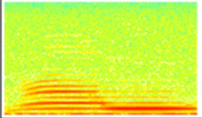 | 0 | 25 | 0 | 0 | 0 | 0 | **25** |
| Waah* | Moggy | 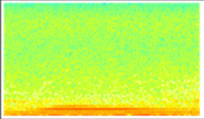 | 0 | 0 | 21 | 0 | 0 | 0 | **21** |
| Waah* | Moggy2 | 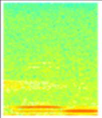 | 0 | 0 | 6 | 0 | 0 | 0 | **6** |
| Waah* | Waah(750Hz) | 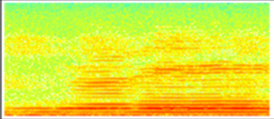 | 0 | 0 | 4 | 0 | 0 | 0 | **4** |
| Waah* | Waah(800hz) | 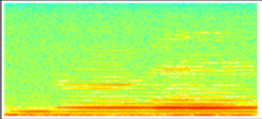 | 0 | 0 | 4 | 0 | 0 | 0 | **4** |

| Broad-scale syllable class | Fine-scale syllable class | Spectrogram | HAU | TAW | TMI | LAI | REP | PKI | Total |
|---|---|---|---|---|---|---|---|---|---|
| Waah* | Waah(850Hz) | 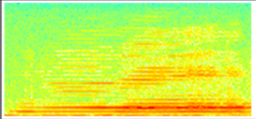 | 0 | 0 | 13 | 0 | 0 | 0 | **13** |
| Waah* | Waah(900Hz) | 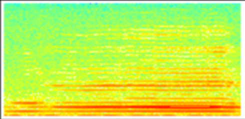 | 0 | 0 | 12 | 0 | 0 | 0 | **12** |
| Waah* | Waah(970Hz) | 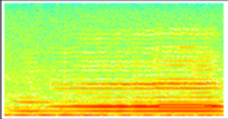 | 0 | 0 | 5 | 0 | 0 | 0 | **5** |
| Waah* | Waah(low+D#) | 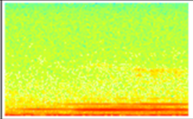 | 0 | 0 | 12 | 0 | 0 | 0 | **12** |
| Waah* | Waah(low+E) | 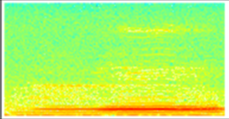 | 0 | 0 | 22 | 0 | 0 | 0 | **22** |
| Waah* | Waah(low+F#) | 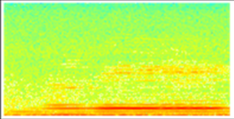 | 0 | 0 | 25 | 0 | 0 | 0 | **25** |
| Waah* | Waah(magpie) | 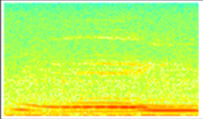 | 0 | 0 | 24 | 0 | 0 | 0 | **24** |
| Waah* | Waah(magpie2) | 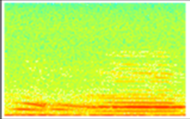 | 0 | 0 | 5 | 0 | 0 | 0 | **5** |
| Waah* | Waah(mid+A#) | 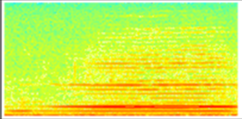 | 0 | 0 | 91 | 0 | 0 | 0 | **91** |

| Broad-scale syllable class | Fine-scale syllable class | Spectrogram | HAU | TAW | TMI | LAI | REP | PKI | Total |
|---|---|---|---|---|---|---|---|---|---|
| Waah* | Waah(mid+A) | 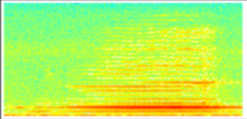 | 0 | 0 | 79 | 0 | 0 | 0 | **79** |
| Waah* | Waah(mid+G#) | 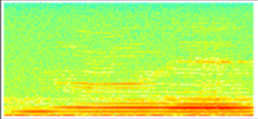 | 0 | 0 | 47 | 0 | 0 | 0 | **47** |
| Waah* | Waah(mid+G) | 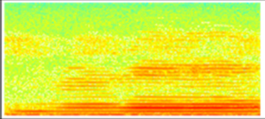 | 0 | 0 | 19 | 0 | 0 | 0 | **19** |
| Waah* | Waah(rough) | 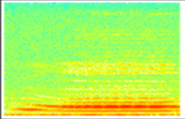 | 0 | 0 | 144 | 0 | 0 | 0 | **144** |
| Waah* | Waah(rough+sansbass) | 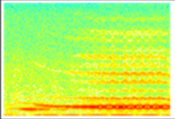 | 0 | 0 | 14 | 0 | 0 | 0 | **14** |
| Waah* | Waah(thin+hi) | 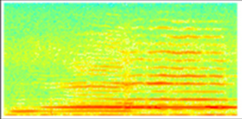 | 0 | 0 | 42 | 0 | 0 | 0 | **42** |
| Waah* | Waah(thin+hi2) | 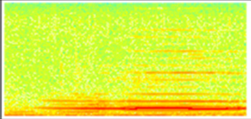 | 0 | 0 | 21 | 0 | 0 | 0 | **21** |
| Waah* | Waah(thin+hi3) | 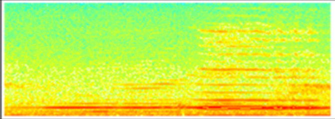 | 0 | 0 | 3 | 0 | 0 | 0 | **3** |
| Waah* | Waah(thin+low) | 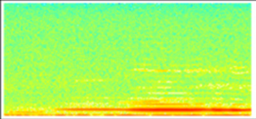 | 0 | 0 | 98 | 0 | 0 | 0 | **98** |

| Broad-scale syllable class | Fine-scale syllable class | Spectrogram | HAU | TAW | TMI | LAI | REP | PKI | Total |
|---|---|---|---|---|---|---|---|---|---|
| Waah* | Waah(thin+med) | 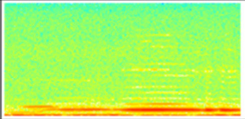 | 0 | 0 | 37 | 0 | 0 | 0 | **37** |
| Waah* | Waah(thin+med2) | 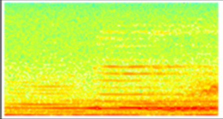 | 0 | 0 | 23 | 0 | 0 | 0 | **23** |
| Waah* | Waah(thin+med3) | 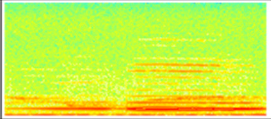 | 0 | 0 | 8 | 0 | 0 | 0 | **8** |
| Warble | Dragon(cleanjump) | 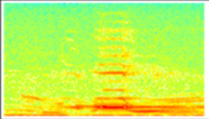 | 0 | 0 | 159 | 0 | 0 | 0 | **159** |
| Warble | Moustache | 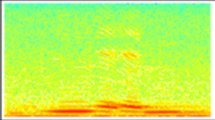 | 0 | 0 | 79 | 0 | 0 | 0 | **79** |
| Warble | Pretrumpet | 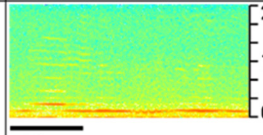  | 0 | 0 | 0 | 0 | 10 | 0 | **10** |
| **Total broad-scale classes: 36** | **Total fine-scale classes: 338** | **Grand totals:** | **551** | **1812** | **4871** | **583** | **3202** | **2317** | **13336** |

### Catalogue of female syllable classes

| Broad-scale syllable class | Fine-scale syllable class | Exemplar spectrogram | HAU | TAW | TMI | LAI | REP | PKI | Total |
|---|---|---|---|---|---|---|---|---|---|
| Alarmy* | Alarmy(short+highbit)* | 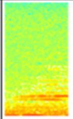 | 0 | 0 | 0 | 0 | 3 | 0 | **3** |
| Alarmy* | Alarmy(short+LBI) | 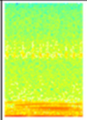 | 4 | 0 | 0 | 0 | 0 | 0 | **4** |
| Alarmy* | Alarmy(short+up)* | 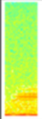 | 0 | 1 | 0 | 0 | 3 | 0 | **4** |
| Alarmy* | Alarmy(trainwhistle2) | 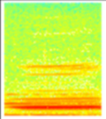 | 0 | 4 | 0 | 0 | 0 | 0 | **4** |
| Alarmy* | Alarmy(twobitCUV+beware)* | 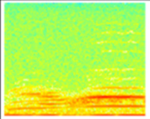 | 0 | 0 | 0 | 0 | 5 | 0 | **5** |
| Alarmy* | Stutter(alarmy)* | 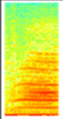 | 0 | 0 | 0 | 2 | 140 | 0 | **142** |
| Alarmy* | Upsqueak(kitten)* | 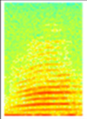 | 0 | 3 | 0 | 0 | 0 | 0 | **3** |
| Chiggle* | Chiggle* | 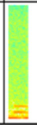 | 0 | 1 | 0 | 0 | 13 | 32 | **46** |

| Broad-scale syllable class | Fine-scale syllable class | Exemplar spectrogram | HAU | TAW | TMI | LAI | REP | PKI | Total |
|---|---|---|---|---|---|---|---|---|---|
| Chirrup | Downcurve(doublevoiced)_Chiup | 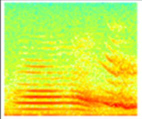 | 0 | 0 | 29 | 0 | 0 | 0 | **29** |
| Chirrup | Stutter_Chiup(decurved) | 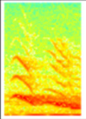 | 0 | 0 | 11 | 0 | 0 | 0 | **11** |
| Chirrup | Stutter_Stutter_Chiup | 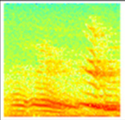 | 0 | 0 | 23 | 0 | 0 | 0 | **23** |
| Chortle* | Killem | 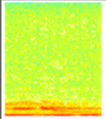 | 0 | 0 | 18 | 0 | 0 | 0 | **18** |
| Chortle* | Killem(2) | 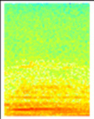 | 0 | 0 | 6 | 0 | 0 | 0 | **6** |
| Chortle* | Nailpolish | 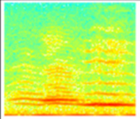 | 0 | 0 | 0 | 0 | 182 | 0 | **182** |
| Chump* | Chiup* | 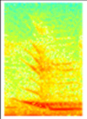 | 0 | 0 | 103 | 0 | 0 | 0 | **103** |
| Cough* | Cough(clickhigh)* | 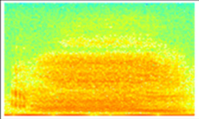 | 10 | 1 | 0 | 0 | 0 | 0 | **11** |
| Cough* | Cough(whip) | 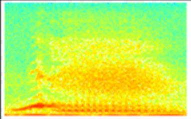 | 0 | 0 | 0 | 0 | 0 | 7 | **7** |

| Broad-scale syllable class | Fine-scale syllable class | Exemplar spectrogram | HAU | TAW | TMI | LAI | REP | PKI | Total |
|---|---|---|---|---|---|---|---|---|---|
| Chirrup | Downcurve(doublevoiced)_Chiup | 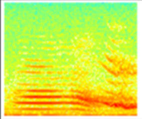 | 0 | 0 | 29 | 0 | 0 | 0 | **29** |
| Chirrup | Stutter_Chiup(decurved) | 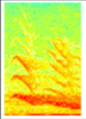 | 0 | 0 | 11 | 0 | 0 | 0 | **11** |
| Chirrup | Stutter_Stutter_Chiup | 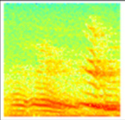 | 0 | 0 | 23 | 0 | 0 | 0 | **23** |
| Chortle* | Killem | 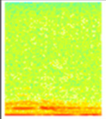 | 0 | 0 | 18 | 0 | 0 | 0 | **18** |
| Chortle* | Killem(2) | 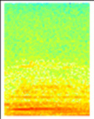 | 0 | 0 | 6 | 0 | 0 | 0 | **6** |
| Chortle* | Nailpolish | 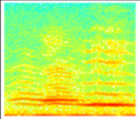 | 0 | 0 | 0 | 0 | 182 | 0 | **182** |
| Chump* | Chiup* | 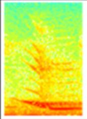 | 0 | 0 | 103 | 0 | 0 | 0 | **103** |
| Cough* | Cough(clickhigh)* | 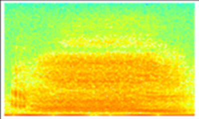 | 10 | 1 | 0 | 0 | 0 | 0 | **11** |
| Cough* | Cough(whip) | 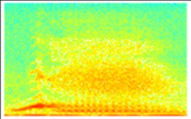 | 0 | 0 | 0 | 0 | 0 | 7 | **7** |

| Broad-scale syllable class | Fine-scale syllable class | Exemplar spectrogram | HAU | TAW | TMI | LAI | REP | PKI | Total |
|---|---|---|---|---|---|---|---|---|---|
| Down* | Down(chip) | 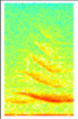 | 0 | 0 | 0 | 0 | 0 | 5 | **5** |
| Down* | Down(CUV1) | 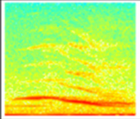 | 0 | 0 | 0 | 0 | 186 | 0 | **186** |
| Down* | Down(CUV2) | 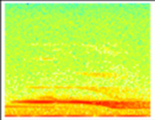 | 0 | 0 | 0 | 0 | 20 | 0 | **20** |
| Down* | Down(LAI2)* | 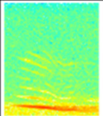 | 0 | 0 | 0 | 7 | 0 | 0 | **7** |
| Down* | Down(LAI3)* | 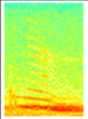 | 0 | 0 | 0 | 53 | 0 | 0 | **53** |
| Down* | Down(LAI4) | 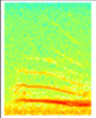 | 0 | 0 | 0 | 15 | 0 | 0 | **15** |
| Down* | Down(LBI1)* | 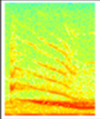 | 6 | 0 | 0 | 0 | 0 | 0 | **6** |
| Down* | Down(notched) | 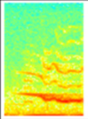 | 0 | 0 | 0 | 0 | 194 | 0 | **194** |
| Down* | Down(PKI+other) | 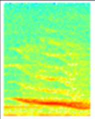 | 0 | 0 | 0 | 0 | 0 | 4 | **4** |

| Broad-scale syllable class | Fine-scale syllable class | Exemplar spectrogram | HAU | TAW | TMI | LAI | REP | PKI | Total |
|---|---|---|---|---|---|---|---|---|---|
| Down* | Down(rough+LBI) | 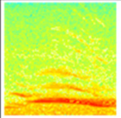 | 8 | 0 | 0 | 0 | 0 | 0 | **8** |
| Down* | Down(slightcurve) | 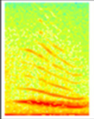 | 0 | 11 | 0 | 0 | 0 | 0 | **11** |
| Down* | Down(slightcurve+click)* | 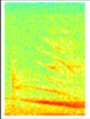 | 10 | 13 | 0 | 0 | 0 | 0 | **23** |
| Down* | Down(slightcurve1)* | 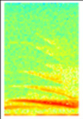 | 12 | 0 | 0 | 0 | 0 | 0 | **12** |
| Down* | Downflick(tiny+TMI) | 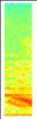 | 0 | 0 | 3 | 0 | 0 | 0 | **3** |
| Down* | Downsweep(burger+LAI) | 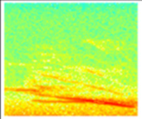 | 0 | 0 | 0 | 4 | 0 | 0 | **4** |
| Down* | Downsweep(flattens) | 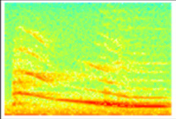 | 0 | 0 | 0 | 14 | 0 | 0 | **14** |
| Down* | Downsweep(roughtosmooth)* | 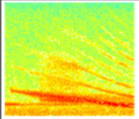 | 0 | 0 | 0 | 174 | 0 | 0 | **174** |
| Down* | Downsweep(roughtosmooth+bigdrop) | 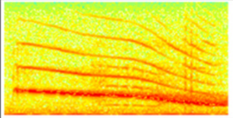 | 0 | 0 | 0 | 22 | 0 | 0 | **22** |

| Broad-scale syllable class | Fine-scale syllable class | Exemplar spectrogram | HAU | TAW | TMI | LAI | REP | PKI | Total |
|---|---|---|---|---|---|---|---|---|---|
| Down_Pipe* | Down_Pipe(A[6]) | 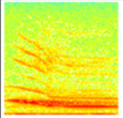 | 1 | 1 | 0 | 1 | 0 | 0 | **3** |
| Down_Pipe* | Down_Pipe(B[6]) | 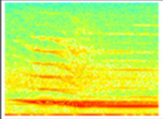 | 0 | 0 | 0 | 0 | 0 | 6 | **6** |
| Down_Pipe* | Down_Pipe(G#[6])* | 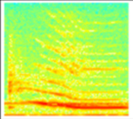 | 16 | 1 | 0 | 0 | 0 | 0 | **17** |
| Down_Pipe* | Down_Pipe(G[6]) | 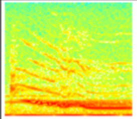 | 43 | 0 | 0 | 0 | 0 | 0 | **43** |
| Down_Pipe* | Down_Pipe(G[6]+)* | 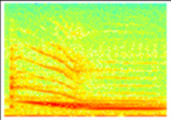 | 25 | 0 | 0 | 0 | 0 | 0 | **25** |
| Down_Pipe* | Downramp_Pipe(D#[7]) | 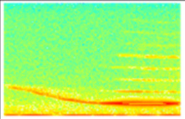 | 0 | 0 | 0 | 0 | 0 | 8 | **8** |
| Down_Pipe* | Downramp_Pipe(D[7]+) | 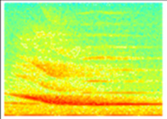 | 0 | 0 | 0 | 0 | 0 | 10 | **10** |
| Downsqueak* | Downcurve(doublevoiced) | 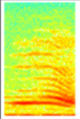 | 0 | 0 | 54 | 0 | 0 | 0 | **54** |
| Downsqueak* | Downramp(alarmy) | 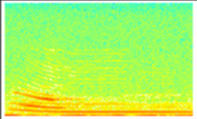 | 5 | 0 | 0 | 0 | 0 | 0 | **5** |

| Broad-scale syllable class | Fine-scale syllable class | Exemplar spectrogram | HAU | TAW | TMI | LAI | REP | PKI | Total |
|---|---|---|---|---|---|---|---|---|---|
| Downsqueak* | Downsqueak | 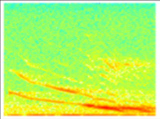 | 13 | 0 | 0 | 0 | 0 | 0 | **13** |
| Downsqueak* | Downsqueak(cleanglass) | 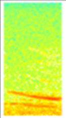 | 8 | 0 | 0 | 0 | 0 | 0 | **8** |
| Downsqueak* | Downsqueak(hook)* | 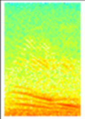 | 0 | 0 | 5 | 0 | 0 | 0 | **5** |
| Downsqueak* | Downsqueak(laser)* | 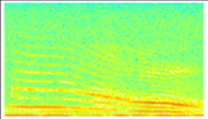 | 0 | 0 | 0 | 0 | 0 | 8 | **8** |
| Downsqueak* | Downsqueak(pew) | 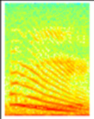 | 0 | 0 | 5 | 0 | 0 | 0 | **5** |
| Downsqueak* | Downsqueak(twopart) | 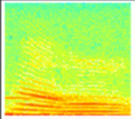 | 3 | 1 | 23 | 0 | 0 | 0 | **27** |
| Downsqueak* | Stutter_Downsqueak(yshape) | 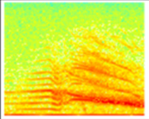 | 0 | 19 | 0 | 0 | 0 | 0 | **19** |
| Downsqueak_Pipe* | Downsqueak(chumpbit)_Pipe(high)* | 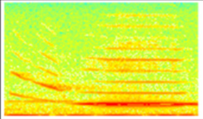 | 3 | 4 | 0 | 0 | 0 | 0 | **7** |
| Flatsqueak* | Pipe(doublevoiced) | 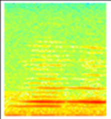 | 5 | 0 | 0 | 0 | 0 | 0 | **5** |

| Broad-scale syllable class | Fine-scale syllable class | Exemplar spectrogram | HAU | TAW | TMI | LAI | REP | PKI | Total |
|---|---|---|---|---|---|---|---|---|---|
| Flatsqueak* | Upsqueak_Flatsqueak(1) | 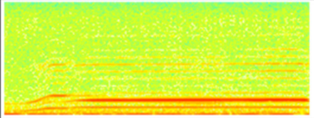 | 0 | 0 | 14 | 0 | 0 | 0 | **14** |
| Flatsqueak* | Upsqueak_Flatsqueak(2) | 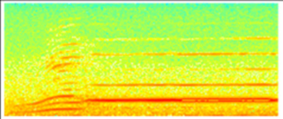 | 0 | 0 | 5 | 0 | 0 | 0 | **5** |
| Flatsqueak* | Upsqueak_Flatsqueak(4) | 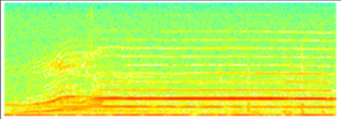 | 0 | 0 | 3 | 0 | 0 | 0 | **3** |
| Peaksqueak* | CUVsqueak* | 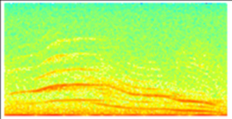 | 0 | 0 | 0 | 0 | 12 | 0 | **12** |
| Peaksqueak* | Peaksqueak(cutting)* | 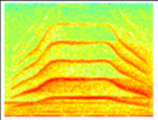 | 0 | 0 | 23 | 0 | 0 | 0 | **23** |
| Peaksqueak* | Peaksqueak(whistley)* | 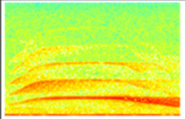 | 0 | 0 | 4 | 0 | 0 | 0 | **4** |
| Peaksqueak* | Pipe(low)_Weeyoo* | 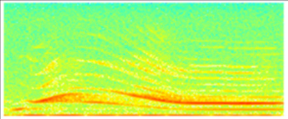 | 0 | 0 | 0 | 0 | 0 | 6 | **6** |
| Peaksqueak* | Pipe_Peak(overlapping) | 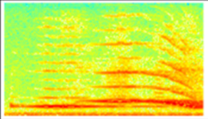 | 0 | 0 | 0 | 0 | 0 | 4 | **4** |
| Peaksqueak* | Upsqueak_Downsqueak | 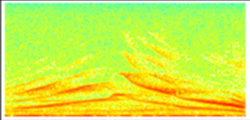 | 6 | 1 | 0 | 0 | 0 | 0 | **7** |

| Broad-scale syllable class | Fine-scale syllable class | Exemplar spectrogram | HAU | TAW | TMI | LAI | REP | PKI | Total |
|---|---|---|---|---|---|---|---|---|---|
| Peaksqueak* | Woah* | 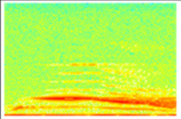 | 0 | 0 | 0 | 0 | 0 | 20 | **20** |
| Pipe* | Pipe(A#[5])* | 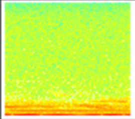 | 3 | 0 | 0 | 0 | 0 | 0 | **3** |
| Pipe* | Pipe(A#[6])* | 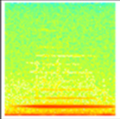 | 16 | 9 | 0 | 11 | 2 | 0 | **38** |
| Pipe* | Pipe(A#[6]+)* | 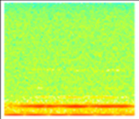 | 30 | 5 | 0 | 5 | 4 | 4 | **48** |
| Pipe* | Pipe(A#[6]+click)* | 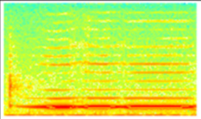 | 8 | 0 | 0 | 0 | 0 | 1 | **9** |
| Pipe* | Pipe(A#[6]+click+) | 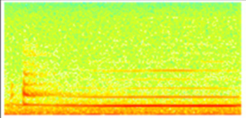 | 8 | 0 | 0 | 0 | 0 | 1 | **9** |
| Pipe* | Pipe(A#[6]bent)* | 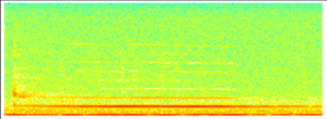 | 5 | 0 | 0 | 0 | 0 | 0 | **5** |
| Pipe* | Pipe(A[6])* | 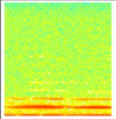 | 3 | 1 | 1 | 1 | 6 | 0 | **12** |
| Pipe* | Pipe(A[6]+)* | 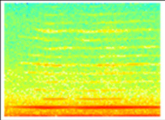 | 6 | 2 | 3 | 1 | 23 | 1 | **36** |

| Broad-scale syllable class | Fine-scale syllable class | Exemplar spectrogram | HAU | TAW | TMI | LAI | REP | PKI | Total |
|---|---|---|---|---|---|---|---|---|---|
| Pipe* | Pipe(B[5])* | 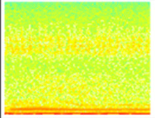 | 3 | 0 | 0 | 0 | 0 | 0 | **3** |
| Pipe* | Pipe(B[6])* | 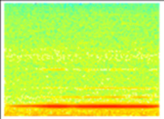 | 14 | 1 | 0 | 2 | 3 | 0 | **20** |
| Pipe* | Pipe(B[6]+)* | 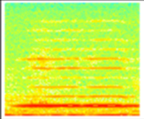 | 5 | 6 | 0 | 1 | 4 | 1 | **17** |
| Pipe* | Pipe(B[6]+click)* | 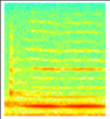 | 12 | 0 | 0 | 0 | 0 | 0 | **12** |
| Pipe* | Pipe(B[7])* | 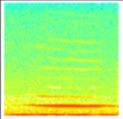 | 1 | 0 | 0 | 0 | 12 | 0 | **13** |
| Pipe* | Pipe(C#[6])* | 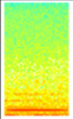 | 2 | 0 | 3 | 0 | 0 | 0 | **5** |
| Pipe* | Pipe(C#[6]+)* | 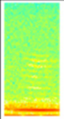 | 2 | 0 | 4 | 0 | 0 | 0 | **6** |
| Pipe* | Pipe(C#[7])* | 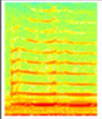 | 3 | 0 | 10 | 0 | 13 | 0 | **26** |
| Pipe* | Pipe(C#[7]+)* | 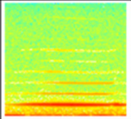 | 20 | 2 | 16 | 0 | 3 | 0 | **41** |

| Broad-scale syllable class | Fine-scale syllable class | Exemplar spectrogram | HAU | TAW | TMI | LAI | REP | PKI | Total |
|---|---|---|---|---|---|---|---|---|---|
| Pipe* | Pipe(C#[7]bent) | 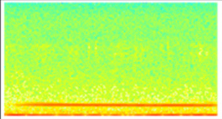 | 1 | 0 | 0 | 4 | 0 | 0 | **5** |
| Pipe* | Pipe(C[7])* | 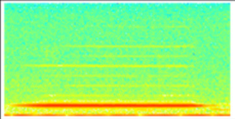 | 4 | 2 | 1 | 0 | 6 | 0 | **13** |
| Pipe* | Pipe(C[7]+)* | 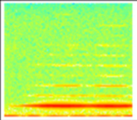 | 1 | 0 | 1 | 4 | 14 | 0 | **20** |
| Pipe* | Pipe(C[7]bent) | 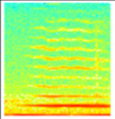 | 1 | 0 | 0 | 0 | 52 | 0 | **53** |
| Pipe* | Pipe(D#[6])* | 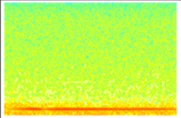 | 8 | 0 | 4 | 0 | 0 | 0 | **12** |
| Pipe* | Pipe(D#[6]+)* | 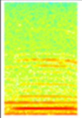 | 2 | 0 | 11 | 0 | 1 | 0 | **14** |
| Pipe* | Pipe(D#[7])* | 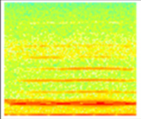 | 5 | 0 | 42 | 1 | 0 | 1 | **49** |
| Pipe* | Pipe(D#[7]+)* | 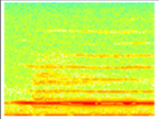 | 3 | 0 | 73 | 6 | 0 | 0 | **82** |
| Pipe* | Pipe(D[6])* | 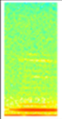 | 4 | 0 | 2 | 0 | 0 | 0 | **6** |

| Broad-scale syllable class | Fine-scale syllable class | Exemplar spectrogram | HAU | TAW | TMI | LAI | REP | PKI | Total |
|---|---|---|---|---|---|---|---|---|---|
| Pipe* | Pipe(D[6]+)* | 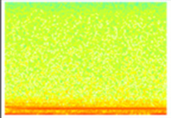 | 5 | 0 | 2 | 0 | 0 | 0 | **7** |
| Pipe* | Pipe(D[7])* | 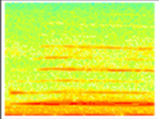 | 18 | 0 | 0 | 2 | 2 | 1 | **23** |
| Pipe* | Pipe(D[7]+)* | 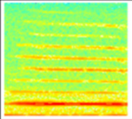 | 3 | 0 | 30 | 0 | 6 | 0 | **39** |
| Pipe* | Pipe(E[6])* | 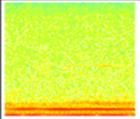 | 4 | 1 | 7 | 0 | 2 | 0 | **14** |
| Pipe* | Pipe(E[6]+)* | 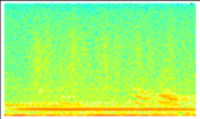 | 1 | 0 | 3 | 0 | 11 | 0 | **15** |
| Pipe* | Pipe(E[6]+ultrashort) | 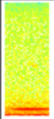 | 3 | 0 | 0 | 0 | 0 | 0 | **3** |
| Pipe* | Pipe(E[7]) | 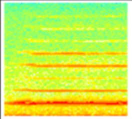 | 1 | 0 | 59 | 9 | 0 | 0 | **69** |
| Pipe* | Pipe(E[7]+) | 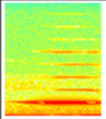 | 1 | 0 | 7 | 21 | 1 | 0 | **30** |
| Pipe* | Pipe(E[7]bent)* | 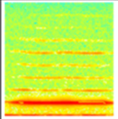 | 3 | 0 | 46 | 0 | 0 | 1 | **50** |

| Broad-scale syllable class | Fine-scale syllable class | Exemplar spectrogram | HAU | TAW | TMI | LAI | REP | PKI | Total |
|---|---|---|---|---|---|---|---|---|---|
| Pipe* | Pipe(F#[6])* | 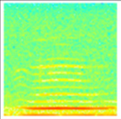 | 18 | 0 | 2 | 0 | 4 | 0 | **24** |
| Pipe* | Pipe(F#[6]+)* | 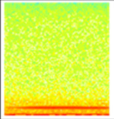 | 8 | 0 | 1 | 0 | 11 | 0 | **20** |
| Pipe* | Pipe(F#[6]+rough) | 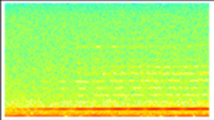 | 0 | 0 | 0 | 0 | 0 | 4 | **4** |
| Pipe* | Pipe(F#[7])* | 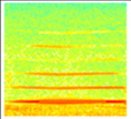 | 2 | 0 | 1 | 1 | 0 | 0 | **4** |
| Pipe* | Pipe(F#[7]+)* | 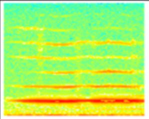 | 2 | 1 | 0 | 1 | 1 | 0 | **5** |
| Pipe* | Pipe(F[6])* | 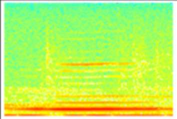 | 10 | 0 | 2 | 0 | 19 | 0 | **31** |
| Pipe* | Pipe(F[6]+)* | 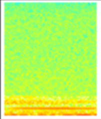 | 0 | 0 | 2 | 0 | 4 | 0 | **6** |
| Pipe* | Pipe(F[6]+ultrashort) | 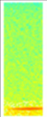 | 3 | 0 | 0 | 0 | 0 | 0 | **3** |
| Pipe* | Pipe(F[6]+whip) | 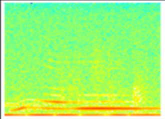 | 0 | 0 | 0 | 0 | 0 | 4 | **4** |

| Broad-scale syllable class | Fine-scale syllable class | Exemplar spectrogram | HAU | TAW | TMI | LAI | REP | PKI | Total |
|---|---|---|---|---|---|---|---|---|---|
| Pipe_Down* | Dinnerplate(1) | 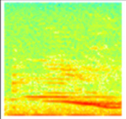 | 21 | 0 | 0 | 0 | 0 | 0 | **21** |
| Pipe_Down* | Dinnerplate(2) | 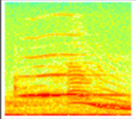 | 4 | 21 | 1 | 0 | 0 | 0 | **26** |
| Pipe_Down* | Dinnerplate(longerplate) | 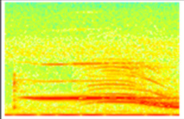 | 3 | 0 | 0 | 0 | 0 | 0 | **3** |
| Pipe_Down* | Peakwhistle(TMI) | 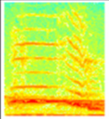 | 0 | 0 | 8 | 0 | 0 | 0 | **8** |
| Pipe_Down* | Pipe(A[7])_Down(LAI) | 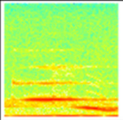 | 0 | 0 | 0 | 10 | 0 | 0 | **10** |
| Pipe_Down* | Pipe(B[6])_Dropoff | 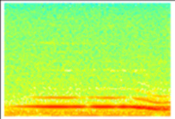 | 0 | 0 | 0 | 8 | 0 | 0 | **8** |
| Pipe_Down* | Pipe(B[6]+)_Dropoff | 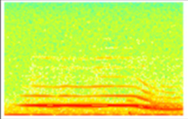 | 0 | 0 | 0 | 9 | 0 | 0 | **9** |
| Pipe_Down* | Pipe(B[7]+)_Down(LAI) | 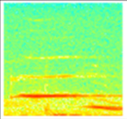 | 0 | 0 | 0 | 3 | 0 | 0 | **3** |
| Pipe_Down* | Pipe(C[7])_Dropoff | 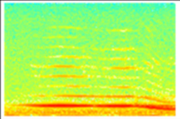 | 0 | 0 | 0 | 45 | 0 | 0 | **45** |

| Broad-scale syllable class | Fine-scale syllable class | Exemplar spectrogram | HAU | TAW | TMI | LAI | REP | PKI | Total |
|---|---|---|---|---|---|---|---|---|---|
| Pipe_Down* | Pipe(C[8])_Down(LAI) | 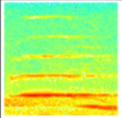 | 0 | 0 | 0 | 5 | 0 | 0 | **5** |
| Pipe_Down* | Pipe(D#[7])_Dropoff | 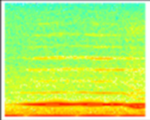 | 1 | 0 | 0 | 0 | 0 | 2 | **3** |
| Pipe_Down* | Pipe(D[7])_Dropoff | 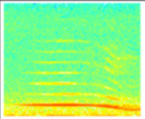 | 0 | 0 | 0 | 5 | 0 | 0 | **5** |
| Pipe_Down* | Pipe(E[7])_Down(LAI) | 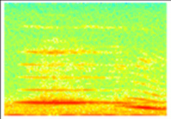 | 0 | 0 | 0 | 15 | 0 | 0 | **15** |
| Pipe_Down* | Pipe(E[7])_Dropoff* | 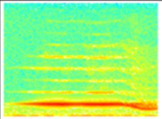 | 0 | 0 | 0 | 0 | 0 | 25 | **25** |
| Pipe_Down* | Pipe(E[7]+)_Dropoff | 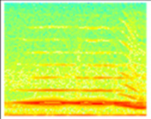 | 0 | 0 | 0 | 0 | 0 | 3 | **3** |
| Pipe_Down* | Pipe(F#[7])_Dropoff | 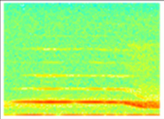 | 2 | 0 | 0 | 0 | 0 | 4 | **6** |
| Pipe_Down* | Pipe(F[7])_Dropoff | 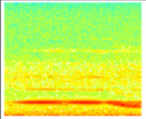 | 0 | 0 | 0 | 0 | 0 | 3 | **3** |
| Pipe_Down* | Pipe(G#[7])_Down | 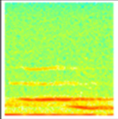 | 0 | 0 | 0 | 19 | 0 | 0 | **19** |

| Broad-scale syllable class | Fine-scale syllable class | Exemplar spectrogram | HAU | TAW | TMI | LAI | REP | PKI | Total |
|---|---|---|---|---|---|---|---|---|---|
| Pipe_Down* | Pipe(G#[7])_Dropoff | 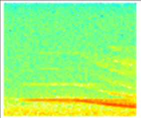 | 0 | 0 | 0 | 3 | 0 | 0 | **3** |
| Pipe_Down* | Pipe(G[7]_Down) | 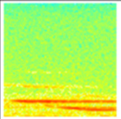 | 0 | 0 | 0 | 15 | 0 | 0 | **15** |
| Pipe_Down* | Pipe(jumpup)_Down(1) | 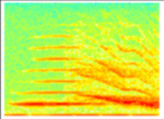 | 0 | 0 | 0 | 0 | 0 | 15 | **15** |
| Pipe_Down* | Pipe(jumpup)_Down(2) | 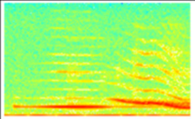 | 0 | 0 | 0 | 0 | 0 | 22 | **22** |
| Pipe_Down* | Pipe(rough)_Tailoff | 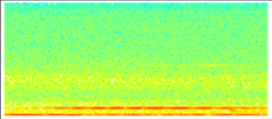 | 0 | 0 | 0 | 0 | 0 | 19 | **19** |
| Pipe_Down_Pipe | Pipe(jumpup)_Down_Pipe(1) | 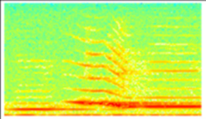 | 0 | 0 | 0 | 0 | 0 | 16 | **16** |
| Pipe_Down_Pipe | Pipe(jumpup)_Down_Pipe(2) | 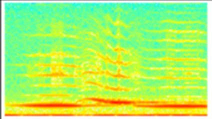 | 0 | 0 | 0 | 0 | 0 | 6 | **6** |
| Pipe_Down_Pipe | Pipe(jumpup)_Down_Pipe(3) | 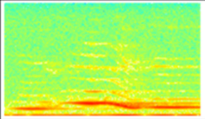 | 0 | 0 | 0 | 0 | 0 | 6 | **6** |
| Pipe_Downsqueak* | Pipe_Downsqueak* | 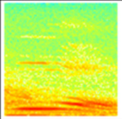 | 0 | 0 | 7 | 0 | 0 | 0 | **7** |

| Broad-scale syllable class | Fine-scale syllable class | Exemplar spectrogram | HAU | TAW | TMI | LAI | REP | PKI | Total |
|---|---|---|---|---|---|---|---|---|---|
| RoughBuzz* | Crescendo(rough)* | 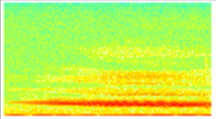 | 0 | 0 | 0 | 5 | 0 | 0 | **5** |
| RoughBuzz* | Down(rough+stepped) | 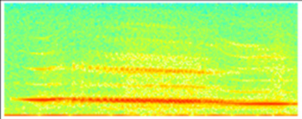 | 0 | 0 | 0 | 0 | 0 | 3 | **3** |
| RoughBuzz* | Whip_Buzz | 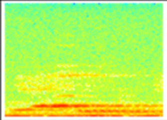 | 0 | 0 | 0 | 0 | 0 | 18 | **18** |
| Shrieky* | Flick(squareroot) | 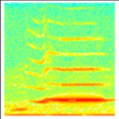 | 0 | 0 | 0 | 0 | 0 | 3 | **3** |
| Shrieky* | Ghh_LBIShriek | 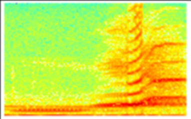 | 18 | 0 | 0 | 0 | 0 | 0 | **18** |
| Shrieky* | LBIShriek | 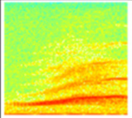 | 12 | 0 | 0 | 0 | 0 | 0 | **12** |
| Shrieky* | Pipe_Upslope_Pipe | 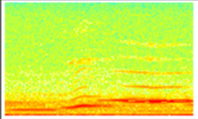 | 1 | 4 | 0 | 0 | 0 | 0 | **5** |
| Stepdown* | DinnerplateFlatArm* | 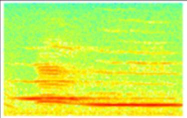 | 15 | 0 | 0 | 0 | 0 | 0 | **15** |
| Stepdown* | DinnerplateFlatArm(rough) | 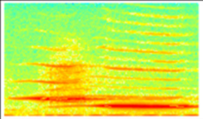 | 8 | 0 | 0 | 0 | 0 | 0 | **8** |

| Broad-scale syllable class | Fine-scale syllable class | Exemplar spectrogram | HAU | TAW | TMI | LAI | REP | PKI | Total |
|---|---|---|---|---|---|---|---|---|---|
| Stepdown* | Down_Click_Stepdown | 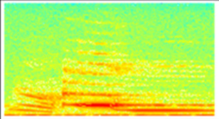 | 0 | 0 | 13 | 0 | 0 | 0 | **13** |
| Stepdown* | Downsqueak_Stepdown(click) | 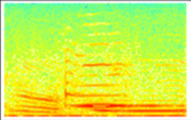 | 0 | 0 | 7 | 0 | 0 | 0 | **7** |
| Stepdown* | Pipe(E[7])_Slide_Pipe(B[6]) | 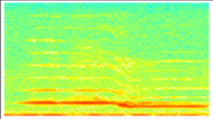 | 0 | 0 | 0 | 0 | 0 | 4 | **4** |
| Stepdown* | PipeOverlap(A#[6]-C[6]) | 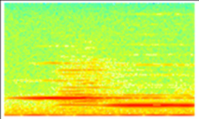 | 6 | 0 | 0 | 0 | 0 | 0 | **6** |
| Stepdown* | Stepdown(A#[7]-A#[6]) | 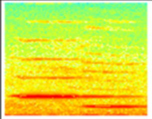 | 0 | 0 | 3 | 0 | 0 | 0 | **3** |
| Stepdown* | Stepdown(A[7]-A[6]) | 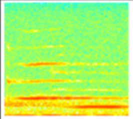 | 0 | 0 | 0 | 5 | 0 | 0 | **5** |
| Stepdown* | Stepdown(A[7]-E[6]) | 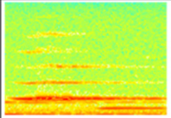 | 3 | 0 | 0 | 0 | 0 | 0 | **3** |
| Stepdown* | Stepdown(A[7]-E[6]+click) | 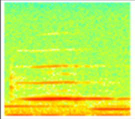 | 5 | 0 | 0 | 0 | 0 | 0 | **5** |
| Stepdown* | Stepdown(C#[7]-C#[6]) | 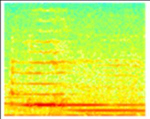 | 0 | 0 | 4 | 0 | 0 | 0 | **4** |

| Broad-scale syllable class | Fine-scale syllable class | Exemplar spectrogram | HAU | TAW | TMI | LAI | REP | PKI | Total |
|---|---|---|---|---|---|---|---|---|---|
| Stepdown* | Stepdown(C#[8]-A[6]) | 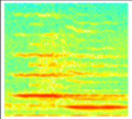 | 0 | 0 | 13 | 0 | 0 | 0 | **13** |
| Stepdown* | Stepdown(C#[8]-D[7]) | 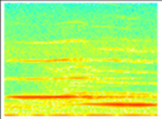 | 3 | 0 | 0 | 0 | 0 | 0 | **3** |
| Stepdown* | Stepdown(C[7]-C[6]) | 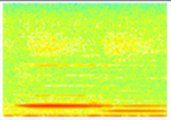 | 0 | 0 | 11 | 0 | 0 | 0 | **11** |
| Stepdown* | Stepdown(C[8]-E[7]) | 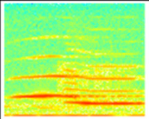 | 4 | 0 | 0 | 1 | 0 | 0 | **5** |
| Stepdown* | Stepdown(D[7]-F[6]) | 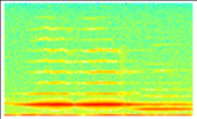 | 0 | 0 | 0 | 0 | 0 | 3 | **3** |
| Stepdown* | Stepdown(F#[7]-A#[6]) | 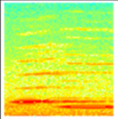 | 1 | 0 | 8 | 0 | 0 | 0 | **9** |
| Stepdown* | Stepdown(F#[7]-A[6]) | 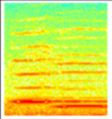 | 0 | 0 | 9 | 0 | 2 | 0 | **11** |
| Stepdown* | Stepdown(F#[7]-B[6]) | 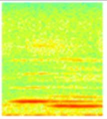 | 2 | 0 | 3 | 0 | 0 | 0 | **5** |
| Stepdown* | Stepdown(F#[7]-E[6]) | 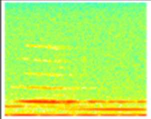 | 0 | 0 | 0 | 6 | 0 | 0 | **6** |

| Broad-scale syllable class | Fine-scale syllable class | Exemplar spectrogram | HAU | TAW | TMI | LAI | REP | PKI | Total |
|---|---|---|---|---|---|---|---|---|---|
| Stepdown* | Stepdown(F#[7]-F#[6]) | 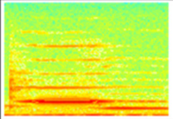 | 0 | 4 | 0 | 0 | 0 | 1 | **5** |
| Stepdown* | Stepdown(F#[8]-D#[6]) | 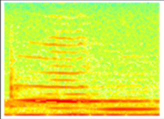 | 0 | 4 | 0 | 0 | 0 | 0 | **4** |
| Stepdown* | Stepdown(F[7]-A#[6]) | 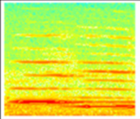 | 0 | 0 | 4 | 1 | 0 | 0 | **5** |
| Stepdown* | Stepdown(F[7]-C#[7]) | 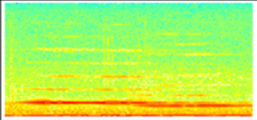 | 0 | 0 | 1 | 0 | 0 | 2 | **3** |
| Stepdown* | Stepdown(G#[7]-D#[6]) | 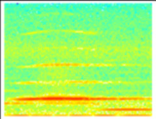 | 6 | 1 | 0 | 0 | 0 | 0 | **7** |
| Stepdown* | Stepdown(G#[7]-D[6]) | 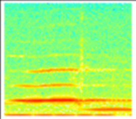 | 4 | 0 | 0 | 0 | 0 | 0 | **4** |
| Stepdown* | Stepdown(G#[7]-E[6]) | 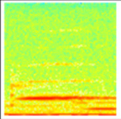 | 10 | 0 | 0 | 0 | 0 | 0 | **10** |
| Stepdown* | Stepdown(G#[7]-G#[6]) | 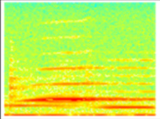 | 0 | 3 | 1 | 0 | 0 | 0 | **4** |
| Stepdown* | Stepdown(G[7]-D#[6])* | 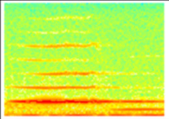 | 4 | 1 | 0 | 0 | 0 | 0 | **5** |

| Broad-scale syllable class | Fine-scale syllable class | Exemplar spectrogram | HAU | TAW | TMI | LAI | REP | PKI | Total |
|---|---|---|---|---|---|---|---|---|---|
| Stepdown* | Upcurve(stepdown)_Pipe(1) | 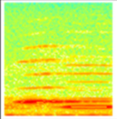 | 0 | 0 | 3 | 0 | 0 | 0 | **3** |
| Stepdown* | Upcurve(stepdown)_Pipe(2) | 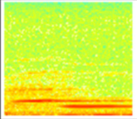 | 0 | 0 | 4 | 0 | 0 | 0 | **4** |
| Stepdown* | Upcurve(stepdown)_Pipe(3) | 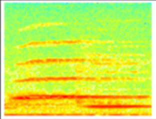 | 0 | 0 | 17 | 0 | 0 | 0 | **17** |
| Stepdown* | Upcurve(stepdown)_Pipe(4) | 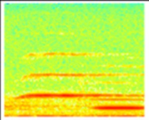 | 0 | 0 | 9 | 0 | 0 | 0 | **9** |
| Stepdown* | Upcurve(stepdown)_Pipe(5) | 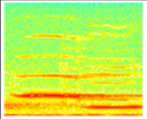 | 0 | 0 | 4 | 0 | 0 | 0 | **4** |
| Stepdown* | Upcurve(stepdown)_Pipe(6) | 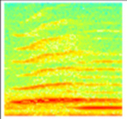 | 0 | 0 | 24 | 0 | 0 | 0 | **24** |
| Stepdown* | Upcurve(stepdown)_Pipe(7) | 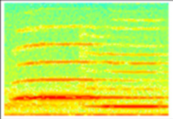 | 0 | 0 | 13 | 0 | 0 | 0 | **13** |
| Stepup* | Pipe(hi+tinystepup1) | 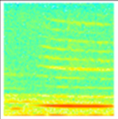 | 0 | 0 | 0 | 0 | 41 | 0 | **41** |
| Stepup* | Pipe(hi+tinystepup3) | 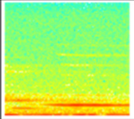 | 0 | 0 | 0 | 0 | 21 | 0 | **21** |

| Broad-scale syllable class | Fine-scale syllable class | Exemplar spectrogram | HAU | TAW | TMI | LAI | REP | PKI | Total |
|---|---|---|---|---|---|---|---|---|---|
| Stepup* | Pipe(hi+tinystepup4) | 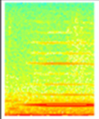 | 0 | 0 | 0 | 0 | 8 | 0 | **8** |
| Stepup* | Pipe(hi+tinystepup5) | 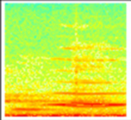 | 0 | 0 | 0 | 0 | 3 | 0 | **3** |
| Stepup* | Stepup(C[7]+-G[7]+) | 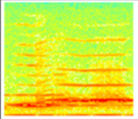 | 0 | 3 | 0 | 0 | 0 | 0 | **3** |
| Stepup* | Stepup(CUV1) | 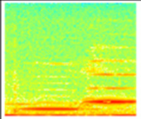 | 0 | 0 | 0 | 0 | 8 | 0 | **8** |
| Stepup* | Stepup(CUV1.5) | 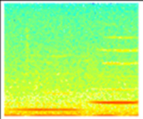 | 0 | 0 | 0 | 0 | 5 | 0 | **5** |
| Stepup* | Stepup(CUV2) | 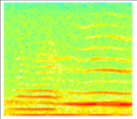 | 0 | 0 | 0 | 0 | 3 | 0 | **3** |
| Stepup* | Stepup(E[6]+-D[7]+) | 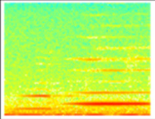 | 0 | 0 | 0 | 0 | 6 | 0 | **6** |
| Stepup* | Stutter_Pipe(D[7]+) | 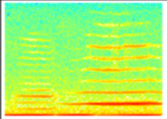 | 0 | 0 | 0 | 0 | 12 | 0 | **12** |
| Stepup_Down* | Pipe(jumpup)_Pipe_Dropoff | 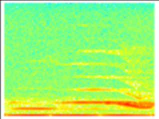 | 0 | 0 | 0 | 0 | 0 | 4 | **4** |

| Broad-scale syllable class | Fine-scale syllable class | Exemplar spectrogram | HAU | TAW | TMI | LAI | REP | PKI | Total |
|---|---|---|---|---|---|---|---|---|---|
| Stutter* | Pipe(D#[6]+ultrashort+) | 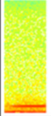 | 4 | 0 | 0 | 0 | 0 | 0 | **4** |
| Stutter* | Stutter* | 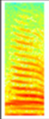 | 211 | 4 | 880 | 26 | 0 | 31 | **1152** |
| Stutter* | Stutter(cheese) | 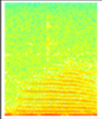 | 0 | 0 | 12 | 0 | 0 | 0 | **12** |
| Stutter* | Stutter(cheet) | 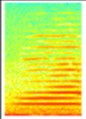 | 0 | 3 | 49 | 0 | 1 | 1 | **54** |
| Stutter* | Stutter(click)* | 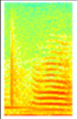 | 0 | 9 | 0 | 0 | 0 | 0 | **9** |
| Stutter* | Stutter(decurved)* | 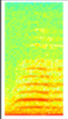 | 0 | 0 | 0 | 0 | 917 | 1 | **918** |
| Stutter* | Stutter(high)* | 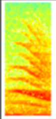 | 17 | 0 | 0 | 0 | 0 | 0 | **17** |
| Stutter* | Stutter(straight) | 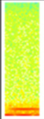 | 3 | 0 | 0 | 0 | 0 | 0 | **3** |
| Stutter* | Stutter(TAW)* | 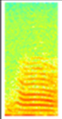 | 0 | 336 | 0 | 0 | 0 | 0 | **336** |

| Broad-scale syllable class | Fine-scale syllable class | Exemplar spectrogram | HAU | TAW | TMI | LAI | REP | PKI | Total |
|---|---|---|---|---|---|---|---|---|---|
| Stutter* | Stutter(wavey) | 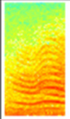 | 0 | 0 | 51 | 0 | 0 | 0 | **51** |
| Stutter* | Stutter(wavey)_Chirp | 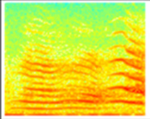 | 0 | 0 | 7 | 0 | 0 | 0 | **7** |
| Stutter* | Stutter_Chirp | 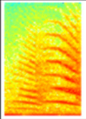 | 0 | 0 | 8 | 0 | 0 | 0 | **8** |
| Stutter(repeats)* | Stutter_Stutter(LBITAW+nownow)* | 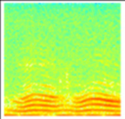 | 6 | 1 | 1 | 0 | 0 | 0 | **8** |
| Stutter_Pipe* | Stutter_Pipe(high+LBI) | 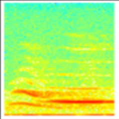 | 15 | 0 | 0 | 0 | 0 | 0 | **15** |
| Stutter_Pipe* | Stutter_Pipe_Down | 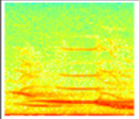 | 6 | 0 | 0 | 0 | 0 | 0 | **6** |
| Trill_Pipe* | Trillpipe | 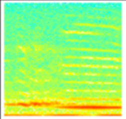 | 0 | 0 | 0 | 0 | 7 | 0 | **7** |
| Upsqueak* | SimUpDown_Upsqueak* | 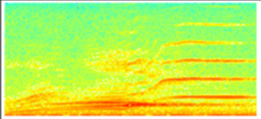 | 0 | 0 | 0 | 0 | 0 | 5 | **5** |
| Upsqueak* | Upsqueak(cleanglass)* | 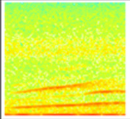 | 10 | 0 | 1 | 0 | 0 | 0 | **11** |

| Broad-scale syllable class | Fine-scale syllable class | Exemplar spectrogram | HAU | TAW | TMI | LAI | REP | PKI | Total |
|---|---|---|---|---|---|---|---|---|---|
| Upsqueak* | Upsqueak(harmonic)* | 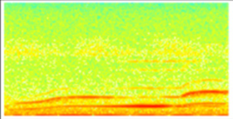 | 5 | 0 | 0 | 0 | 0 | 0 | **5** |
| Upsqueak* | Upsqueak(twopeaks) | 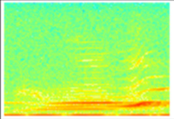 | 0 | 0 | 0 | 0 | 0 | 6 | **6** |
| Waah* | Cat(TMI1) | 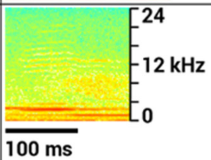  | 0 | 0 | 3 | 0 | 0 | 0 | **3** |
| **Total broad-scale classes: 27** | **Total fine-scale classes: 218** | **Grand totals:** | **854** | **491** | **1873** | **546** | **2087** | **334** | **6185** |
