## Supplementary figures and images for "Sexually distinct song cultures in a songbird metapopulation"

### Figure S1

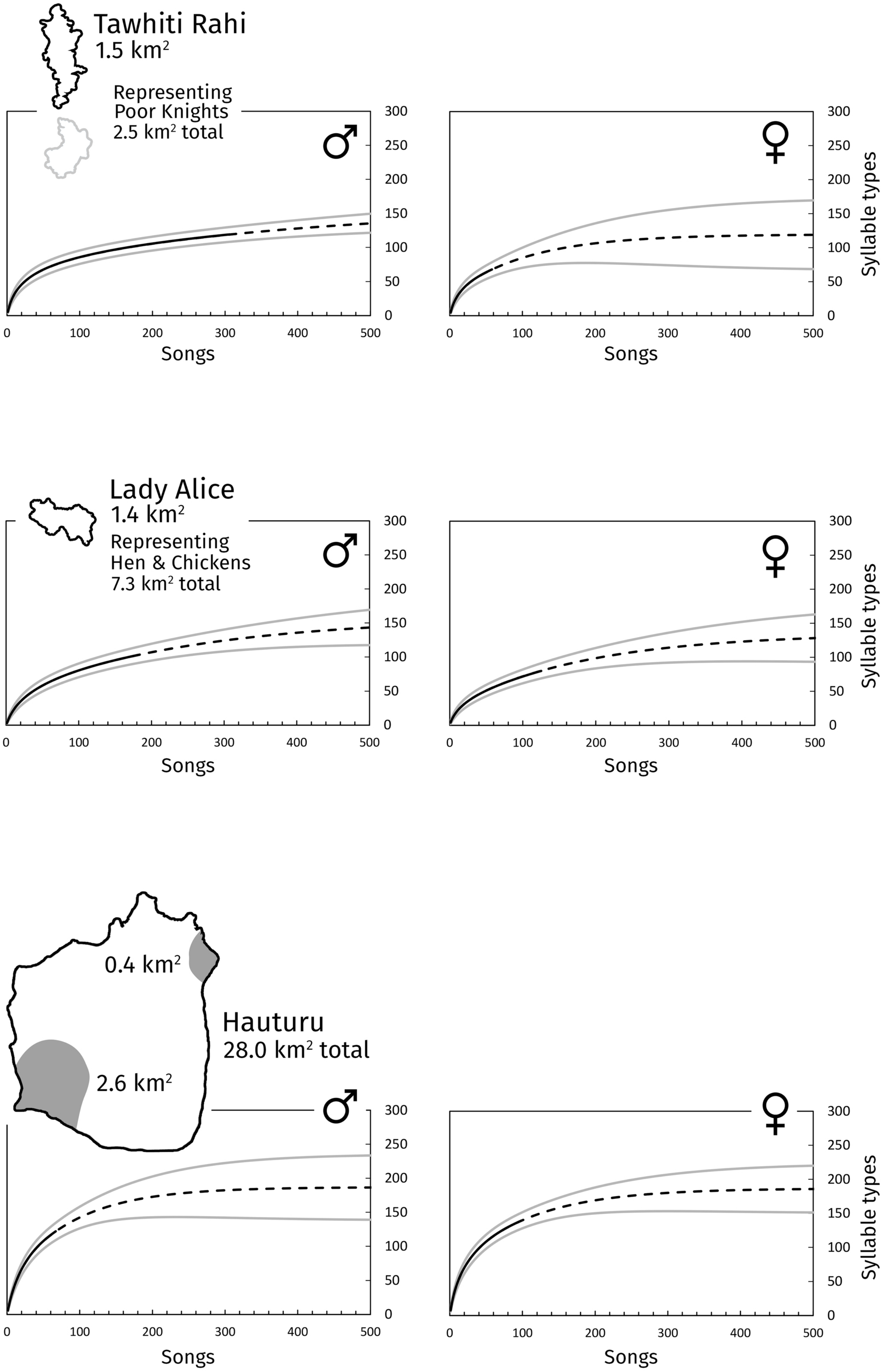

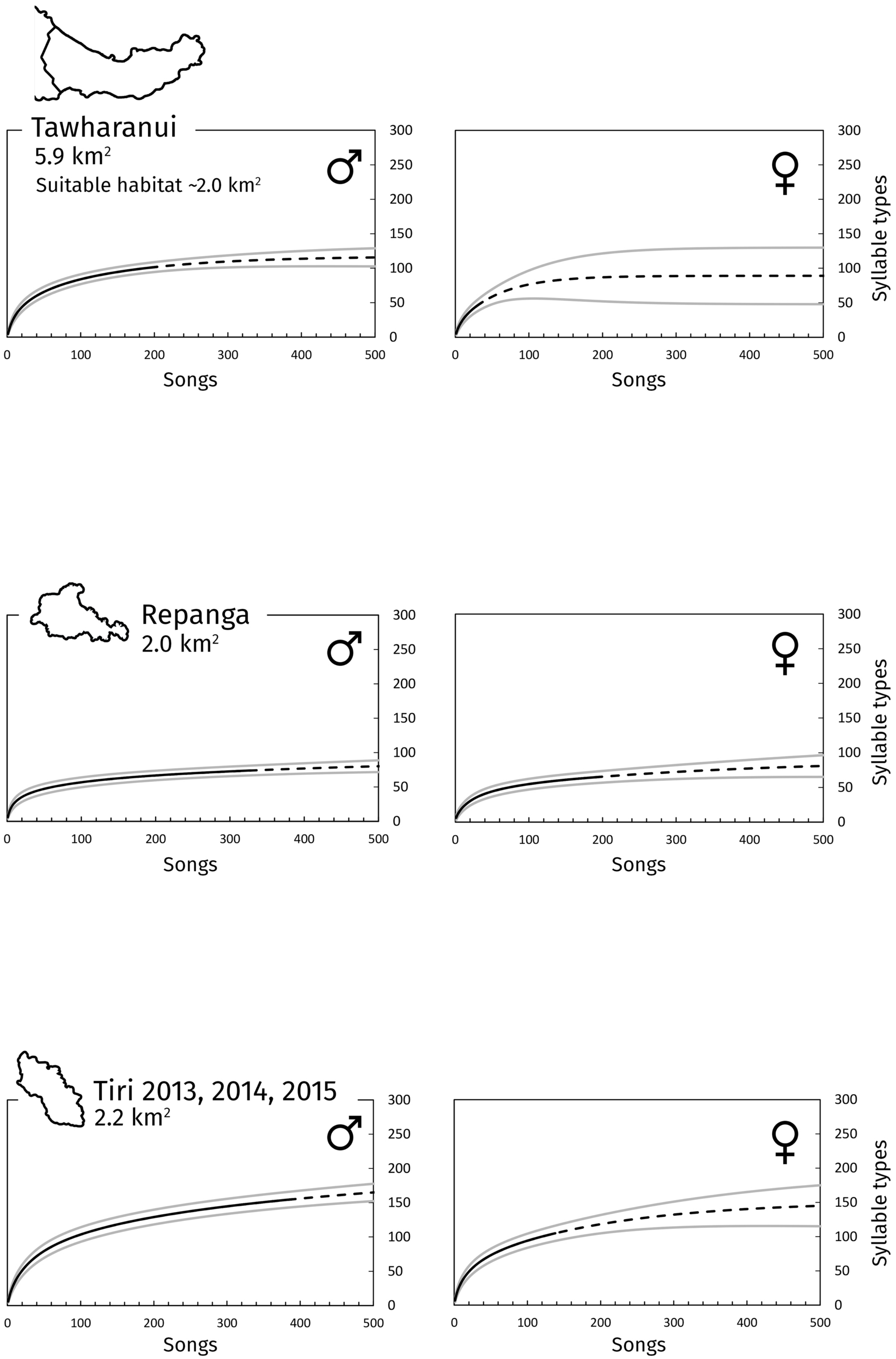
