## Supplementary material for "Sexually distinct song cultures in a songbird metapopulation": Table S2

#### Syllable types sung by both males and females at each site

The list for each site corresponds to the male-female bar overlap regions in Figure 2 of the main article. Scalebars are provided.

| Tawhiti Rahi | |
|---|---|
| **Fine-scale syllable class** | **Spectrogram** |
| Chiggle | 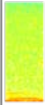 |
| Downsqueak(laser) | 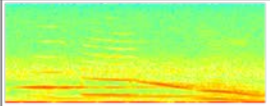 |
| Pipe(low)_Weeyoo | 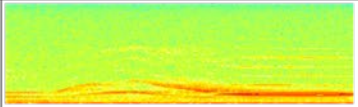 |
| SimUpDown_Upsqueak | 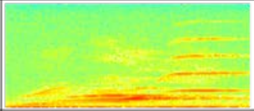 |
| Stutter | 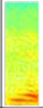 |
| Woah | 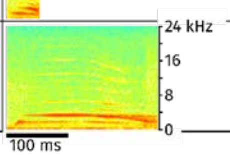 |

| Lady Alice Island | |
|---|---|
| **Fine-scale syllable class** | **Spectrogram** |
| Crescendo(rough) | 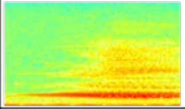 |
| Down(LAI2) | 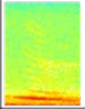 |
| Down(LAI3) | 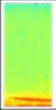 |
| Downsweep(roughtosmooth) | 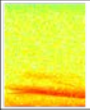 |
| Pipe(A#[6]) | 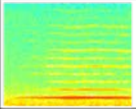 |
| Pipe(A#[6]+) | 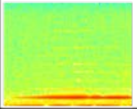 |
| Stutter | 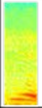 |

#### Hauturu

| Fine-scale syllable class | Spectrogram | Fine-scale syllable class | Spectrogram |
|---|---|---|---|
| Cough(clickhigh) | 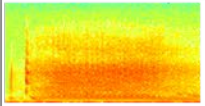 | Pipe(A#[6]bent) | 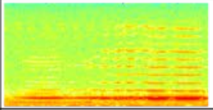 |
| DinnerplateFlatArm | 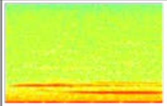 | Pipe(A[6]) | 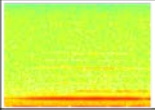 |
| Down(LBI1) | 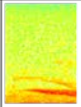 | Pipe(A[6]+) | 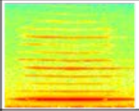 |
| Down(slightcurve+click) | 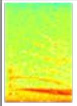 | Pipe(B[6]) | 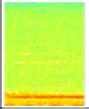 |
| Down(slightcurve1) | 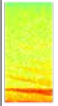 | Pipe(D#[6]) | 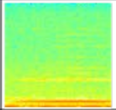 |
| Down_Pipe(G#[6]) | 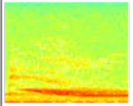 | Pipe(D[6]) | 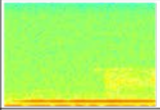 |
| Down_Pipe(G[6]+) | 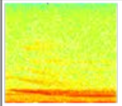 | Pipe(D[6]+) | 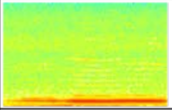 |
| Pipe(A#[6]) | 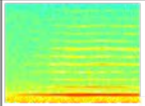 | Pipe(D[7]) | 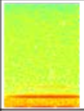 |
| Pipe(A#[6]+) | 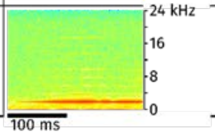  | Pipe(D[7]+) | 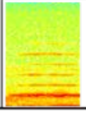 |

#### **Hauturu** *(cont.)*

| Fine-scale syllable class | Spectrogram |
|---|---|
| Pipe(E[6]) | 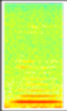 |
| Pipe(F#[6]) | 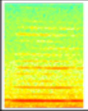 |
| Pipe(F#[6]+) | 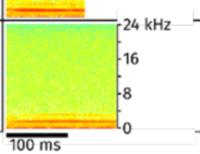  |

| Fine-scale syllable class | Spectrogram |
|---|---|
| Pipe(F[6]) | 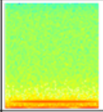 |
| Stepdown(G[7]-D#[6]) | 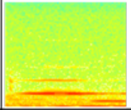 |
| Stutter | 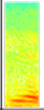 |
| Stutter(high) | 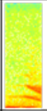 |
| Stutter_Stutter(LBITAW+nownow) | 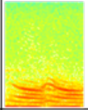 |
| Upsqueak(cleanglass) | 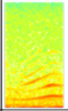 |
| Upsqueak(harmonic) | 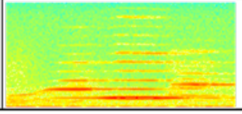 |

### Repanga

| Fine-scale syllable class | Spectrogram | Fine-scale syllable class | Spectrogram |
|---|---|---|---|
| Alarmy(short+highbit) | 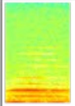 | Pipe(C[7]+) | 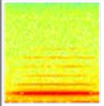 |
| Alarmy(short+up) | 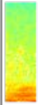 | Pipe(E[6]+) | 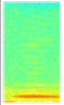 |
| Alarmy(twobitCUV+beware) | 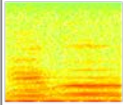 | Pipe(F#[6]) | 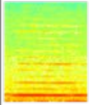 |
| Chiggle | 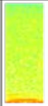 | Pipe(F#[6]+) | 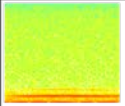 |
| CUVsqueak | 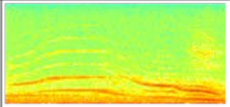 | Pipe(F[6]) | 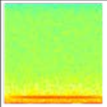 |
| Pipe(A#[6]+) | 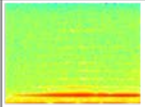 | Pipe(F[6]+) | 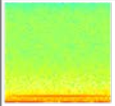 |
| Pipe(A[6]+) | 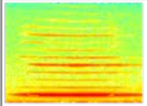 | Pipe(F[7]+) | 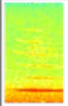 |
| Pipe(B[7]) | 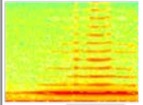 | Stutter(alarmy) | 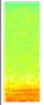 |
| Pipe(C#[7]) | 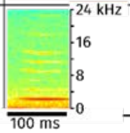  | Stutter(decurved) | 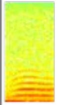 |

| Tawharanui | |
|---|---|
| **Fine-scale syllable class** | **Spectrogram** |
| Downsqueak(chumpbit)_Pipe(high) | 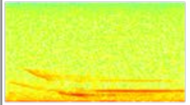 |
| Pipe(A#[6]) | 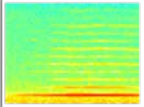 |
| Pipe(B[6]+) | 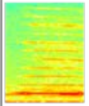 |
| Pipe(G[6]+) | 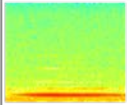 |
| Stutter | 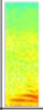 |
| Stutter(click) | 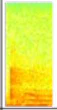 |
| Stutter(TAW) | 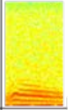 |
| Upsqueak(kitten) | 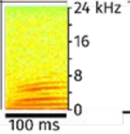  |

| Tiri 2013 | |
|---|---|
| **Fine-scale syllable class** | **Spectrogram** |
| Chiup | 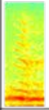 |
| Peaksqueak(cutting) | 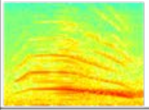 |
| Peaksqueak(whistley) | 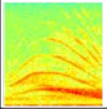 |
| Pipe(A[6]+) | 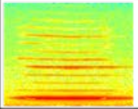 |
| Pipe(C#[6]) | 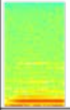 |
| Pipe_Downsqueak | 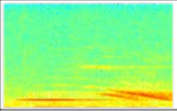 |
| Stutter | 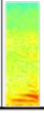 |

| Tiri 2014 | |
|---|---|
| **Fine-scale syllable class** | **Spectrogram** |
| Downsqueak(hook) | 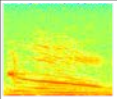 |
| Peaksqueak(cutting) | 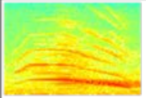 |
| Pipe(A[6]+) | 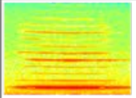 |
| Pipe(C#[6]+) | 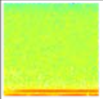 |
| Pipe(D#[6]) | 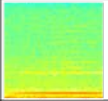 |
| Pipe_Downsqueak | 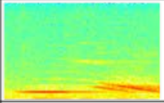 |
| Stutter | 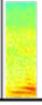 |

| Tiri 2015 | |
|---|---|
| **Fine-scale syllable class** | **Spectrogram** |
| Chiup | 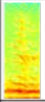 |
| Downsqueak(hook) | 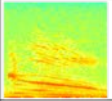 |
| Peaksqueak(cutting) | 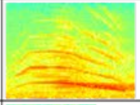 |
| Peaksqueak(whistley) | 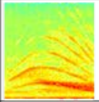 |
| Pipe(C#[6]) | 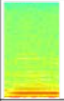 |
| Pipe(D#[6]) | 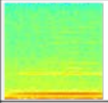 |
| Pipe(E[6]) | 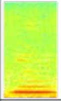 |
| Pipe(E[6]+) | 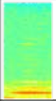 |
| Pipe_Downsqueak | 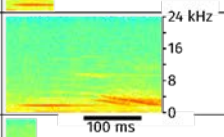  |
| Stutter | 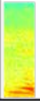 |

#### Tiri 2013, 2014, 2015

| Fine-scale syllable class | Spectrogram | Fine-scale syllable class | Spectrogram |
|---|---|---|---|
| Chiup | 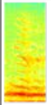 | Pipe(D#[6]) | 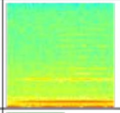 |
| Downsqueak(hook) | 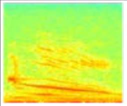 | Pipe(E[6]) | 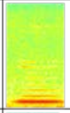 |
| Peaksqueak(cutting) | 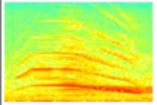 | Pipe(E[6]+) | 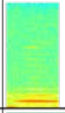 |
| Peaksqueak(whistley) | 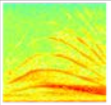 | Pipe(G#[6]) | 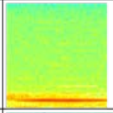 |
| Pipe(A[6]+) | 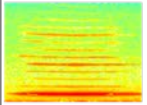 | Pipe_Downsqueak | 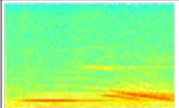 |
| Pipe(C#[6]) | 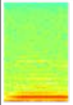 | Stutter | 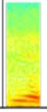 |
| Pipe(C#[6]+) | 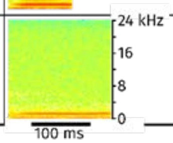  | | |
